# Nitro Reduction-Based RNA Control and Ultrafast Release

**DOI:** 10.64898/2026.02.28.708682

**Authors:** Yiran Zhao, Senfeng Zhang, Junsong Guo, Tuan-Khoa Kha, Song Chen, Chunyi Hu, Ru-Yi Zhu

**Affiliations:** Department of Chemistry, National University of Singapore, 117543 Singapore; Department of Biological Sciences, National University of Singapore, 117559 Singapore

**Keywords:** Post-synthetic modification, nitro reduction, RNA caging

## Abstract

RNA protection and controlled release are critical for both fundamental research and therapeutic applications, yet the development of simple, efficient, and reversible post-synthetic RNA modification strategies remains a significant challenge. Here, we introduce a straightforward approach based on ribose 2′-hydroxyl acylation with a nitro-functionalized carbamate, which acts as a redox-responsive center. This modification selectively inhibits native RNA function while remaining inert to endogenous biogenic reductants and common reducing agents used in biological assays. Upon treatment with low millimolar concentrations of the THDB– BIPY reducing pair, RNA function is rapidly restored on a minute timescale. This methodology is broadly applicable across diverse RNA classes and functional contexts, including synthetic RNA oligomers, fluorogenic RNA aptamers, single-guide RNAs in CRISPR–Cas9 gene-editing systems, and mRNAs in living cells for translation control and RNAi-mediated gene silencing. These results demonstrate the general utility of this approach as a chemically controllable functional switch, providing a versatile toolkit for temporal regulation of RNA activity in both research and biotechnological applications.

---

RNA is a versatile and essential class of biological macro-molecules that plays central roles in protein translation^1, 2^, regulation of gene expression^3–5^, and diverse aspects of cellular function.^6^ Its structural and functional plasticity underlies broad opportunities in both fundamental research and clinical applications. The development of RNA-based therapeutics—including small interfering RNA (siRNA), antisense oligonucleotide (ASO), microRNA, messenger RNA (mRNA), CRISPR-associated single guide RNA (sgRNA)^7–10^—has advanced rapidly, culminating in multiple agents approved by the U.S. Food and Drug Administration (FDA).^11^ To enhance stability, reduce immunogenicity, and improve pharmacological performance, extensive efforts have been devoted to chemical modification of RNA, targeting the nucleobase (e.g., pseudouridine)^12, 13^, the ribose moiety (e.g., 2’-OMe)^14^ and the phosphate backbone (e.g., phosphorothioate linkage).^15, 16^

In the context of biomolecular chemical modification, most established strategies rely on stable, irreversible transformations that enhance molecular stability or efficacy, or enable probing and labeling of specific biological interactions.^17–19^ In contrast, reversible modification introduces functionally disruptive moieties that act as temporary “locks”, attenuating native activity. Upon exposure to exogenous or endogenous stimuli, selective bond cleavage restores the unmodified biomolecule.^20–23^ This stimulus-responsive reversibility expands opportunities for precision medicine and has been broadly applied in peptide- and protein-based therapeutics.^24, 25^ In comparison, analogous strategies for RNA remain underdeveloped. The discovery of highly responsive, bioorthogonal, and RNA-compatible reversible chemistries is therefore of substantial importance for advancing nucleic acid–based therapeutic modalities.

Although photoresponsive^26^, enzyme-triggered^27^ and click chemistry-induced^28^ strategies for reversible RNA protection have been reported, their broader implementation has been constrained by the need for synthetically elaborate modified monomers and their applicability primarily to short RNA oligomers^29^. In contrast, post-synthetic RNA modification enables installation of functional groups without complex phosphoramidite preparation, offering clear advantages in cost, scalability, and accessibility. Leveraging the high nucleophilicity of the ribose 2′-hydroxyl (2’-OH) group^30–32^, *N*-acyl imidazole–mediated 2’-OH acylation has emerged as a powerful platform for RNA probing and engineering, including structural mapping^33–35^ and proximity labeling.^36–38^ Such acylation markedly attenuates RNA function by sterically perturbing secondary structure formation and protein recognition relative to the native state. These features render 2′-O-acylation an attractive scaffold for the development of stimulus-responsive, control–release strategies.

Pioneering work by Kool and co-workers introduced cleavable RNA acylation systems reversible either through phosphine-mediated Staudinger reduction^39^ or blue-light–induced Norrish type II photodeprotection^40^, representing a significant advance in post-synthetic reversible RNA acylation. However, these approaches exhibit notable limitations: photoresponsive acylation does not achieve complete substrate conversion, whereas phosphine-triggered deprotection proceeds relatively slowly and relies on toxic P(III) reagents, restricting applications in living systems. More recently, the Zhu group reported thiol-responsive reversible RNA acylation applicable to diverse RNA substrates, demonstrating its practical utility.^41, 42^ This strategy exploits endogenous thiols, such as glutathione (GSH), enabling spontaneous deprotection under conditions of elevated intracellular GSH levels. Despite these advances, the development of an orthogonal reversible acylation platform featuring rapid installation and deprotection kinetics, triggered by exogenous stimuli and unaffected by endogenous metabolites or biomolecules, remains an unmet challenge for RNA manipulation in biological contexts.

The nitro group, characterized by a high oxidation state of nitrogen, is readily reduced to the corresponding aniline when appended to aromatic systems, a transformation widely exploited in prodrug design.^43, 44^ Nitroaromatic caging motifs responsive to nitroreductase (NTR)—an enzyme upregulated in hypoxic tumor cells but minimally active under normoxic conditions—have been extensively investigated for selective small-molecule activation.^45–47^ Despite promising preclinical results, clinical translation of such systems has thus far remained limited.^48^

In parallel, a new class of diboron–bipyridine reducing systems has emerged as an efficient and chemoselective platform for nitroaromatic reduction under mild conditions.^49, 50^ Notably, the Chu group demonstrated that water-soluble tetrahydroxydiboron (THDB, **A**) in combination with 4,4′-bipyridine (BIPY, **B**) enables effective prodrug activation in cultured cells and *in vivo* mouse models, highlighting its biocompatibility and functional robustness in biological environments.^51^ Inspired by the reductive conversion of *para*-nitroaryl groups to anilines, we envisioned coupling this transformation with a self-immolative linker architecture bearing a benzyl leaving group (**Figure 1A**). Reduction of the nitro group to the corresponding amine at the *para* position is expected to initiate a 1,6-elimination cascade, driven by entropic gain associated with carbon dioxide extrusion from a carbonate linkage. The rapid kinetics of this decarboxylative fragmentation suggest its suitability as a trigger for an ultrafast, post-synthetic, control–release RNA modification platform.

**FIGURE 1.**
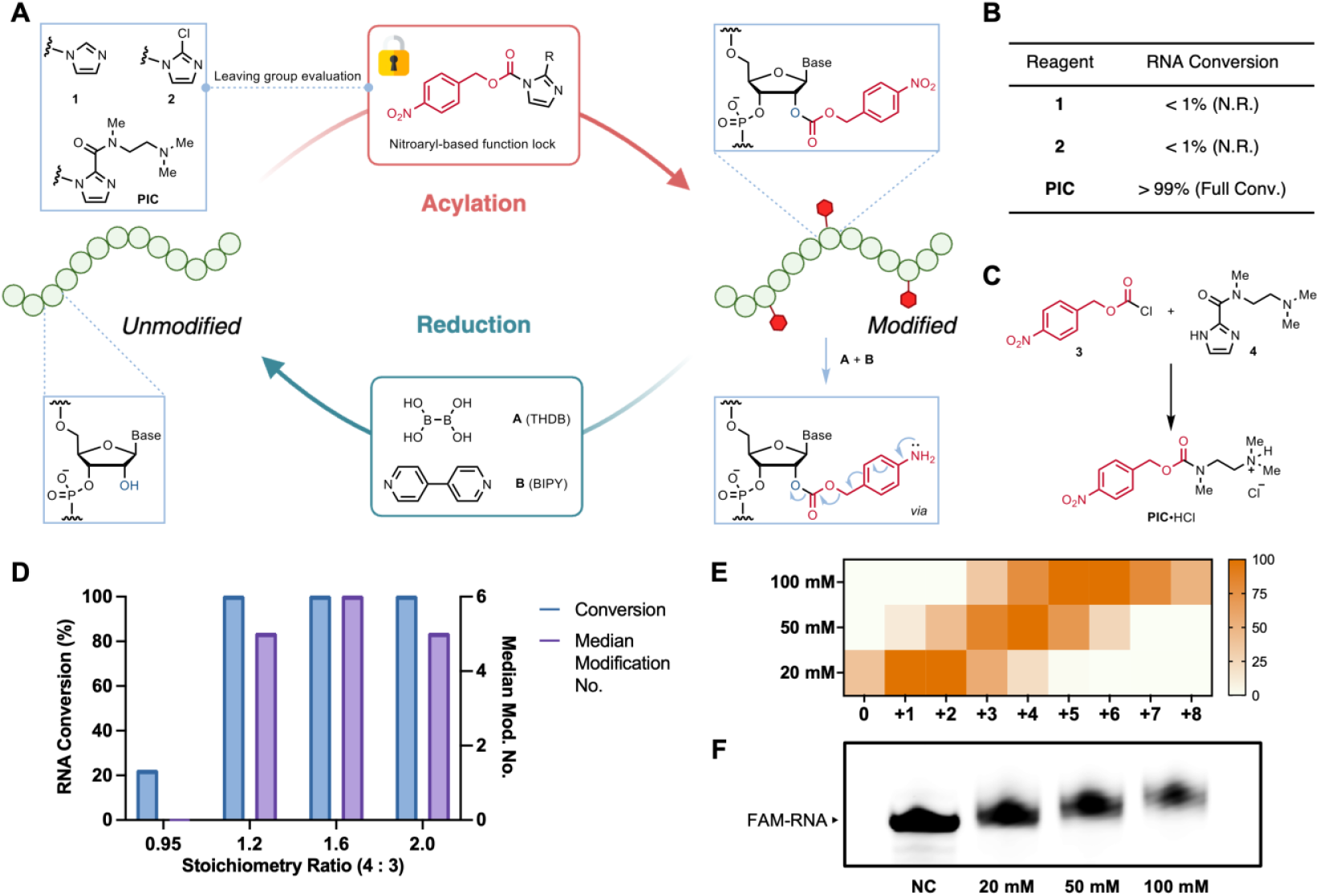
Chemical discovery of p-nitrobenzyl imidazole-2-amide carbamate (**PIC**) and concept validation of nitro-reduction responsive RNA acylation strategy. A) Schematic design and proposed mechanism of RNA 2’-OH acylation with nitrobenzyl-based acylating agents and reductive release by tetrahydroxydiboron (THDB) and 4,4’-bipyridine (BIPY). Structure of leaving group moieties in **1, 2**, and **PIC** is symbolized by imidazole substituted with R group. B) Performance screening of nitro-based acylation agent with different leaving group on model 18-mer FAM-RNA. C) Synthetic approach of **PIC** preparation by chloroformate **3** and imidazole-2-amide **4**. D) Performance screening of **PIC** chloroformate-imidazole ratio on 18-mer FAM-RNA. Conversion in B) and D) was evaluated from MALDI-TOF analysis. E) Heat map of 18-mer FAM-RNA modification number (by integration of MALDI peak area and with largest peak normalized to 100%) when treated with 20, 50 and 100 mM **PIC** and F) gel electrophoresis visualization of acylated FAM-RNA. All acylation experiment was conducted in buffer-free, RNase-free water at room temperature (25 °C) for 120 min, with final *x*(DMSO) = 20%.

## RESULTS AND DISCUSSION

### Nitro reduction-based RNA acylation and release

To validate the proposed control–release strategy, we first designed a redox-responsive acylating agent and evaluated suitable leaving groups for installing a carbonate linkage onto RNA oligomers. The reagent architecture comprises three functional elements: (i) a redox-responsive nitro group, (ii) a self-immolative benzyl alcohol scaffold capable of 1,6-elimination, and (iii) a leaving group to enable RNA acylation. Guided by precedent in leaving-group design, we synthesized candidate reagents bearing imidazole and 2-chloroimidazole as leaving groups (**1** and **2**, respectively; **Figure 1A**), each prepared as 500 mM stock solutions in anhydrous DMSO. An 18-nucleotide (nt) FAM-labeled RNA oligonucleotide was selected as a model substrate to assess acylation efficiency. However, treatment with 100 mM of **1** or **2** at room temperature resulted in minimal conversion, as determined by MALDI-TOF mass spectrometry (**Figure 1B** and **S1**). Substantial precipitation was observed upon addition of the acylating agents to the aqueous RNA solution, suggesting poor solubility and consequently low effective concentrations in contact with the oligonucleotide.

To address this limitation, we adopted a water-soluble imidazole-based leaving-group strategy^52^ by incorporating a tertiary amine side chain to enhance hydrophilicity. This hydrophilic imidazole was subsequently incorporated into the *para*-nitro-benzyl carbamate scaffold to generate *para*-nitrobenzyl imidazole-2-amide carbamate (**PIC**) (**Figure 1C**). Addition of **PIC** to aqueous RNA solutions produced markedly less precipitation and enabled near-complete conversion of the RNA substrate (**Figure 1B**). Notably, the increased polarity and electrophilicity of **PIC** rendered it incompatible with reverse-phase chromatography following synthesis. We therefore optimized the preparation by tuning the stoichiometry between imidazole-2-amide (**4**) and *para*-nitrobenzyl chloroformate (**3**) (**Figure 1C**). Unexpectedly, no acylation occurred when the reagents were combined in a 1:1 ratio during synthesis, whereas use of 20–100% excess imidazole **4** afforded full conversion. Optimal results were obtained with 1.6 equiv of imidazole **4**, yielding a median acylation number of +6 on the 18-mer RNA at 100 mM reagent concentration and room temperature (**Figure 1D** and **S2**). This optimized batch was subsequently used for further studies. Concentration-dependent RNA modification was then evaluated by treating the oligonucleotide with 20, 50, and 100 mM **PIC**, resulting in median modification levels of +1, +3, and +6, respectively, as determined by MALDI-TOF analysis (**Figure 1E** and **S3**). These trends were corroborated by 20% denaturing polyacrylamide gel electrophoresis (PAGE), which showed progressive mobility shifts consistent with increasing degrees of acylation (**Figure 1F**).

Following purification of the acylated RNA, we evaluated reductive deprotection under biologically relevant conditions using both enzymatic and chemical approaches. Treatment with recombinant *E. coli* nitroreductase (NTR), even at 500 μg/mL in the presence of 5 mM NADH in 1×PBS, resulted in incomplete reduction after 24 h at 37 °C (**Figure S4**). The limited enzymatic turnover is likely attributable to steric shielding of the nitroaryl moiety by RNA. In contrast, exposure to a chemical reducing system comprising 4 mM **A** and 1 mM **B**, premixed in a 4:1 stoichiometric ratio as previously described^51^, generated a highly reactive intermediate^49^ (**Figure 2A**) and led to rapid, quantitative restoration of the native RNA within 20 min (**Figure 2B**). Complete conversion was confirmed by MALDI-TOF mass spectrometry (**Figure 2C**), demonstrating efficient RNA release at low millimolar concentrations of small-molecule reductants on a remarkably short timescale.

**FIGURE 2.**
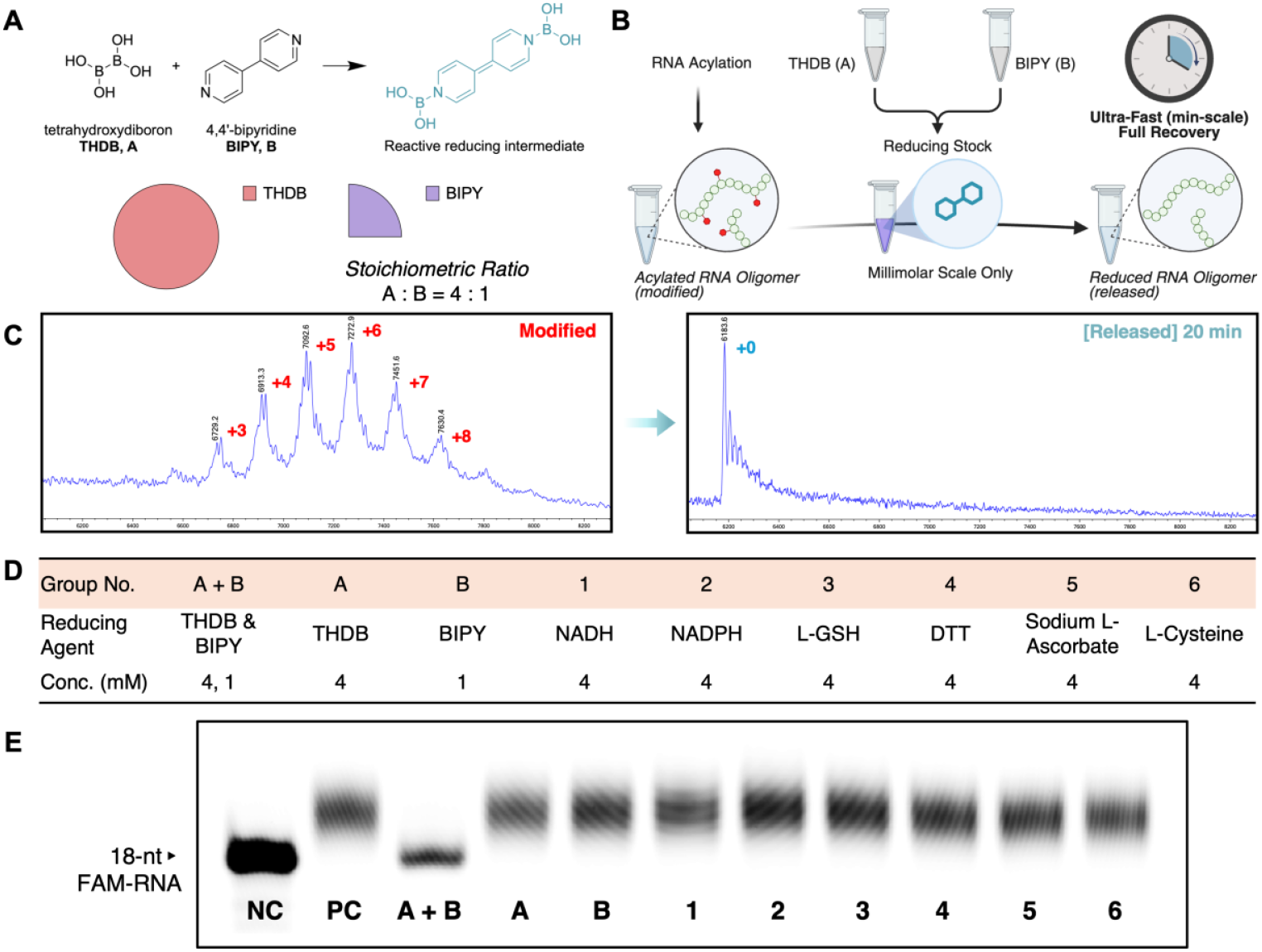
Ultra-fast RNA reduction-elimination when treated with reducing agent. A) Stoichiometry and structure of THDB and BIPY and chemical structure of proposed intermediate. B) Illustration of in vitro RNA reduction when treated with 4 mM THDB and 1 mM BIPY premix. C) MALDI-TOF of 100 mM acylated RNA before and after reduction with 4 mM THDB and 1 mM BIPY for 20 min in 1X PBS, 37 °C. Red and blue numbers stand for modification numbers on one single RNA molecule (calculated by mass difference, each modification account for around +180). D) Reducing agent type and concentration in modification reduction bioorthogonal validation and E) denatured polyacrylamide electrophoresis analysis of the RNA after incubation for 20 min in 1X PBS, 37 °C. NC, unmodified FAM-RNA; PC, 100 mM **PIC** acylated FAM-RNA. All other labels symbolize different reductant treatment.

To assess orthogonality and stability, **PIC**-modified RNA was incubated with **A** or **B** individually, as well as with biologically relevant reductants—including dihydronicotinamide derivatives, thiols, and ascorbate—under identical conditions (except for **B** concentration) for 20 min (**Figure 2D**). No detectable deprotection was observed with any single component or endogenous reducing agent. Only the combined **A**/**B** system effected complete restoration of the native RNA (**Figure 2E**). These results establish **PIC**-mediated RNA acylation as a bioorthogonal modification platform that is selectively and rapidly reversed by the **A**/**B** reducing pair, while remaining inert to endogenous metabolites and common biochemical assay reagents.

### *In vitro* RNA functional control-recovery

Having established that **PIC** acylation of a model 39-nt RNA disrupts Watson–Crick base pairing—as evidenced by melting temperature analysis—and that this effect is fully reversible upon reductive triggering (**Figure S5**), we next evaluated functional control and recovery in a structured RNA system. Because **PIC** installation perturbs both intra- and intermolecular interactions, we anticipated that RNA function would be suppressed upon acylation and restored only after treatment with the **A**/**B** reducing system. Fluorogenic RNA aptamers provide a convenient platform to assess this hypothesis. These structured RNAs bind cognate small-molecule dyes and, upon proper folding, induce strong fluorescence. Steric perturbation of the aptamer scaffold is expected to disrupt folding and abolish fluorescence, whereas reductive removal of the acyl groups should restore native structure and signal (**Figure 3A**). We selected the 104-nt green fluorogenic aptamer F30-Pepper (**Figure 3B**), which binds HBC530 (**Figure 3C**),^53, 54^ as a model system. Following *in vitro* transcription, the aptamer was treated with 20, 50, or 100 mM **PIC** (DMSO alone as control), purified by spin column, quantified, and incubated in 1×folding buffer containing 5 μM HBC530. Fluorescence imaging revealed a clear, concentration-dependent decrease in fluorescence intensity upon acylation (**Figure 3D**). Reductive treatment of the modified aptamer with 4 mM **A** and 1 mM **B** led to time-dependent fluorescence recovery, as visualized at 1, 5, 20, and 60 min (**Figure 3E**). Quantitative analysis using a microplate reader, based on full emission spectra (**Figure S6**), confirmed both the concentration-dependent suppression of fluorescence (**Figure 3F**) and the rapid kinetics of functional restoration (**Figure 3G**). At 100 mM **PIC**, fluorescence was reduced to <10% of the native signal, whereas exposure to the **A**/**B** system restored full fluorescence intensity within 5 min. These results demonstrate efficient, reversible shutdown and ultrafast recovery of RNA function through **PIC**-mediated acylation and chemically triggered deprotection.

**FIGURE 3.**
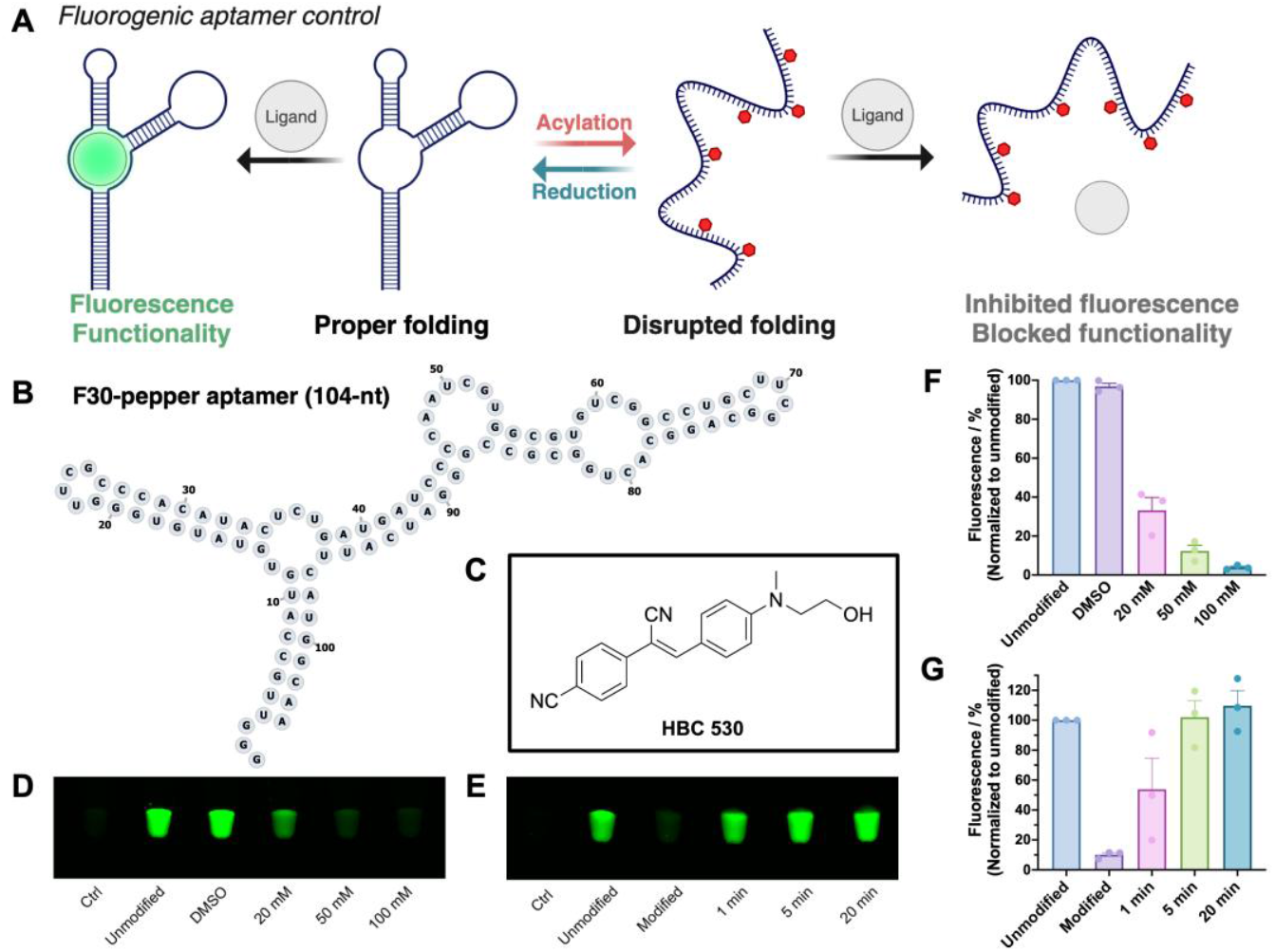
F30-pepper aptamer fluorescence control. A) Demonstration of RNA control-release system application on fluorogenic RNA aptamers. B) Sequence and structural information of 104-nt F30-pepper aptamer. C) Chemical structure of pepper aptamer ligand, HBC530. D) and E) Visualization of F30-pepper aptamer fluorescence with respect to D) **PIC** concentration and E) kinetics of A/B-mediated reductive recovery, on gel scanner. Data are representatives of three replicate experiments. F) and G) Quantification of samples in D) and E) by fluorescence emission spectra on microplate reader. Acylation of D) and F) was conducted in buffer-free water at room temperature (25 °C) for 120 min. Reduction of E) and G) was conducted with 4 / 1 mM THDB and BIPY in 1X PBS, 37 °C. Ctrl: HBC530 only. DMSO: mock treatment of 20% DMSO. Modified, 50 mM PIC acylated aptamer. Data are presented as the mean ± SEM of n = 3 independent experiments.

As a foundational gene-editing platform, CRISPR (Clustered Regularly Interspaced Short Palindromic Repeats) technology, adapted from bacterial adaptive immune systems, has been extensively engineered and deployed across diverse biological settings.^55, 56^ In the canonical system, the endonuclease Cas9 is directed by a single guide RNA (sgRNA), which comprises a Cas9-binding scaffold and a DNA-targeting spacer sequence. Through Watson–Crick base pairing between the spacer and the complementary genomic DNA strand, Cas9 is positioned to introduce site-specific double-strand breaks via its nuclease-active catalytic domains. We reasoned that installation of the control–release modification onto sgRNA would disrupt both Cas9 binding and target DNA hybridization through steric interference from acylation, thereby suppressing gene-editing activity. Subsequent treatment with the **A**/**B** reducing system should restore sgRNA structure and function *in situ*. To test this hypothesis, we selected a 4 kb double-stranded DNA substrate as the target and designed a corresponding 131-nt sgRNA based on the *Neisseria meningitidis* Cas9 (Nme1Cas9) platform (**Figure 4A**). Cas9 protein was preincubated with either native or **PIC**-modified sgRNA prior to exposure to the DNA substrate. Experimental groups included an untreated DNA negative control (NC) and groups 1–6 corresponding to native sgRNA, **PIC**-modified sgRNA, and modified sgRNA subjected to varying concentrations of reducing agents (**Figure 4B**). Cleavage efficiency was quantified by agarose gel electrophoresis, comparing the intensity of uncleaved DNA to the combined intensities of truncated products (**Figure 4C**). Direct treatment of the reaction mixture with **A** and **B** for 20 min resulted in a concentration-dependent restoration of nuclease activity. At the highest reducing-agent concentration (group 6), editing efficiency was fully recovered and statistically indistinguishable from that of native sgRNA (group 1). In contrast, treatment with **A** or **B** alone failed to reinstate sgRNA function (**Figure S7**). Collectively, these results demonstrate that **PIC**-mediated reversible acylation is highly compatible with the CRISPR–Cas9 system, enabling temporally controlled and rapidly activatable gene-editing activity. This strategy holds promise as a chemically gated toolkit for precision genome engineering.

**FIGURE 4.**
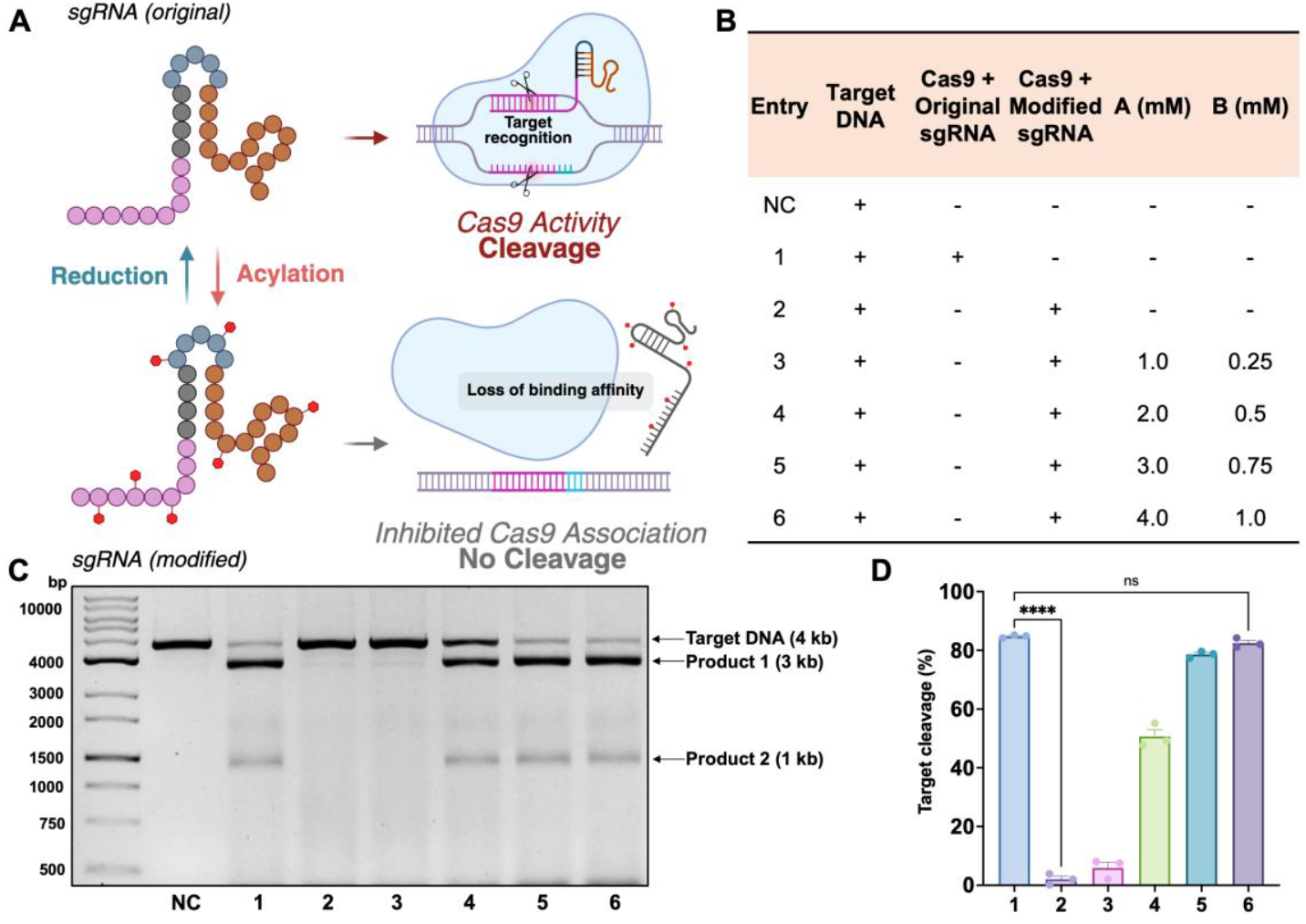
CRISPR-Cas9 system functional control by sgRNA modification and release. A) Schematic illustration of downstream effect of original and modified sgRNA on Cas9 enzyme binding and activity. B) Outline of different entries components of sgRNA function experiment in C) and D). All experiments were treated one-pot, and the reduction was incubated for 20 minutes at room temperature (25 °C). C) Gel electrophoresis visualization of target dsDNA cleavage (with reference to target and product fragment DNA length). Data are representatives of three replicate experiments. D) Quantification of relative percentage of target cleavage based on band intensity percentage of gel electrophoresis. Data are presented as the mean ± SEM of n = 3 independent experiments.

### *In cellulo* mRNA function recovery

Having established proof-of-concept on short-to medium-length (<200 nt) RNAs *in vitro*, we next extended the strategy to longer transcripts. Messenger RNAs encoding fluorescent proteins, such as enhanced green fluorescent protein (EGFP), with typically around 1000 nt in length, offer convenient readouts for visualization and quantification both *in vitro* and *in cellulo* (via lipid-mediated transfection). We therefore selected EGFP mRNA (mEGFP) as a model substrate to evaluate inhibition–recovery of long-RNA function. Translation of mRNA requires coordinated intermolecular interactions, including binding of poly(A)-binding protein (PABP) to the poly(A) tail and assembly of the eIF4F initiation complex.^2, 57^ We anticipated that **PIC**-mediated acylation would sterically disrupt these interactions and suppress translation, with restoration achievable upon treatment with the **A**/**B** reducing pair (**Figure 5A**). Initial treatment of mEGFP with 50 mM **PIC** led to poor recovery during purification (<30% yield), likely reflecting substantial changes in RNA hydrophobicity that compromised spin-column retention. Because 20 mM **PIC** afforded incomplete modification (**Figure 1E**), we selected 35 mM **PIC** for subsequent *in vitro* studies. Following reductive treatment and repurification, equal amounts of native, acylated, or released mEGFP were subjected to a wheat germ extract translation system, and EGFP fluorescence was quantified at 530 nm. As expected, acylation reduced translation to <10% of native levels. Reductive treatment with 4 mM **A** and 1 mM **B** partially restored fluorescence to ~55% of the original signal (**Figure 5B**). To assess RNA integrity, unmodified, modified, and released mEGFP samples were analyzed on 1% agarose gels. No smearing or additional bands were observed (**Figure S8**), indicating the absence of detectable degradation during either acylation or reduction. The incomplete functional recovery may arise from imperfect refolding of the long transcript after acylation or from residual modification persisting on a subset of sites. Nonetheless, fluorescence in the released group was significantly increased relative to the acylated group (ANOVA, p < 0.0001), supporting further evaluation in cellular systems. For *in cellulo* studies, HeLa cells were transfected with native or **PIC**-modified mEGFP. Six hours post-transfection, cells were treated with 4 mM **A** and 1 mM **B** for 120 min and then incubated for an additional 16 h to allow protein expression. Initially, **A** and **B** stocks were prepared in DMSO, but this resulted in minimal recovery—likely due to the mildly oxidative nature of DMSO interfering with the diboron reducing system. Switching to aqueous stock solutions resolved this issue (**Figure S9**). Under optimized conditions, confocal microscopy revealed strong suppression of EGFP expression in the modified group, with clear restoration upon treatment with the reducing small molecules under cellular conditions (**Figure 5C**). For quantitative analysis, cells were transfected with unmodified or 25, 30, or 35 mM **PIC**-modified mEGFP, with or without **A**/**B** treatment, and analyzed by flow cytometry after 16 h (**Figure 5D**). EGFP expression decreased in a **PIC** concentration–dependent manner, whereas **A**/**B** treatment shifted the fluorescence distribution toward higher intensity within the EGFP-positive population. Analysis of median GFP fluorescence (**Figure 5E**) confirmed substantial restoration of expression by the reducing pair. Collectively, these results demonstrate that **PIC**-mediated reversible acylation can modulate the function of long mRNA transcripts and enable chemically triggered reactivation *in vitro* and in living cells. Although further optimization may be required to achieve higher recovery efficiencies for long RNAs, this platform establishes a foundation for temporally controlled regulation of mRNA translation.

**FIGURE 5.**
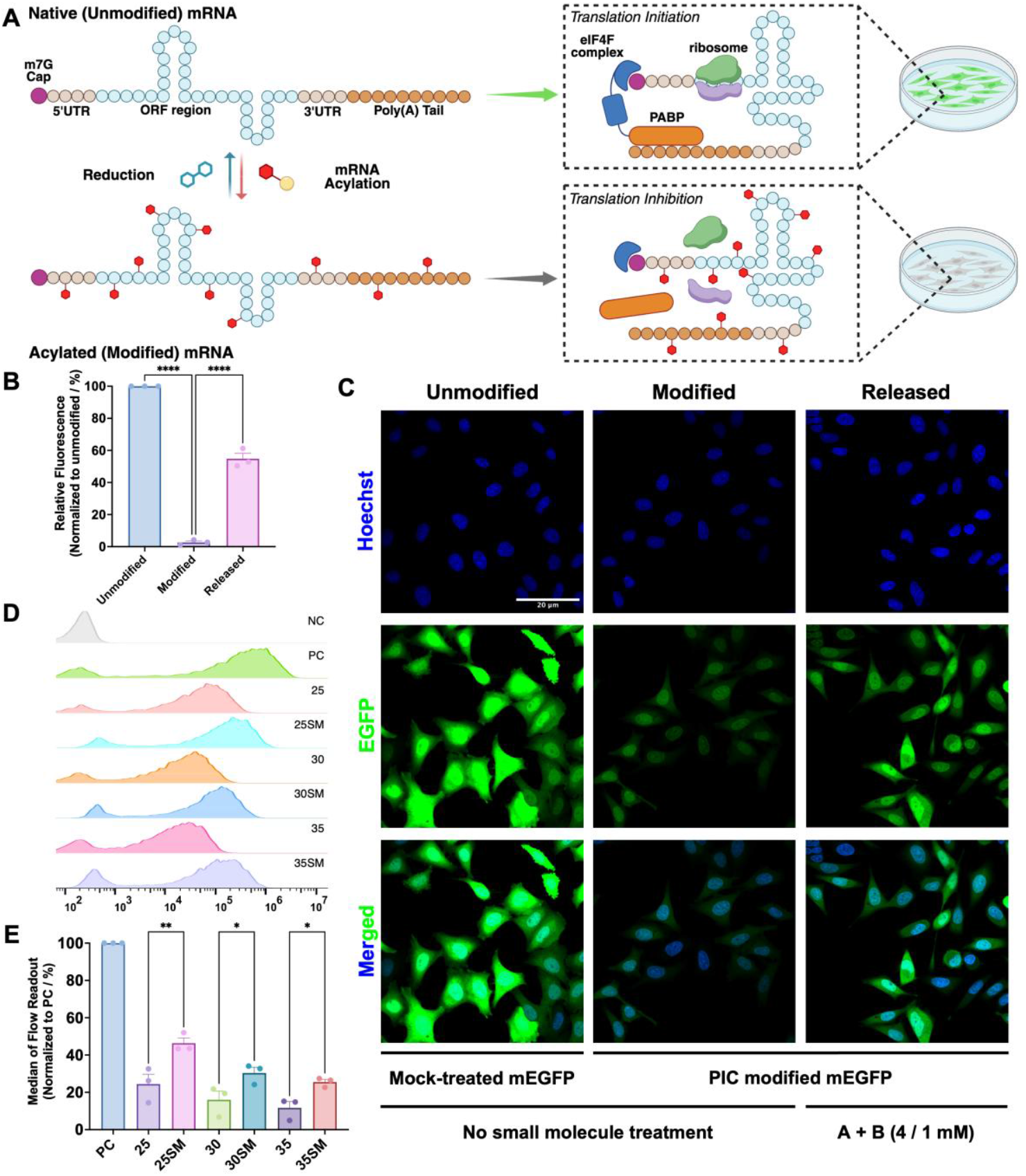
mRNA *in vitro* and *in cellulo* control-recovery. A) Mechanism of translation initiation and inhibition for unmodified (or released) and modified mRNA. B) Intensity of EGFP mRNA (mEGFP) *in vitro* translation. For unmodified (mock treated with 7% DMSO only), modified (acylated with 35 mM **PIC**, 7% DMSO) and released (reduction with 4 mM A and 1 mM B for 20 min, 37 °C) groups, mRNA was purified with spin column, and an equal amount of respective mRNA was injected into *in vitro* translation system and incubated at room temperature (25 °C) for 120 min. C) Confocal microscopy of HeLa cells transfected with unmodified (mock-treated) or **PIC** modified (35 mM) mEGFP for 6 h and treated with blank treatment (1X PBS) or mixture of 4 mM A and 1 mM B in 1X PBS for 120 min, then further cultured for 16 h. Images are representative of three independent experiments. D) Flow cytometry histogram demonstration and E) quantification of median cell fluorescence of unmodified, modified and *in cellulo* released mEGFP signal. NC, cells without any transfection. PC, cells transfected with unmodified mEGFP. 25, 30, 35, cells transfected with mEGFP modified with 25, 30, 35 mM **PIC**. SM, cells released with reducing small molecule treatment after transfection of the respective **PIC**-modified mEGFP. Data are presented as the mean ± SEM of n = 3 independent experiments.

### Caged short hairpin RNA as controllable RNAi strategy

Encouraged by the successful control–release of RNA function in living cells, we next explored the applicability of this platform as a chemical caging strategy for gene-specific silencing. RNA interference (RNAi) relies on cellular processing of synthetic RNA into double-stranded duplexes that guide formation of the RNA-induced silencing complex (RISC), ultimately leading to RNase-mediated degradation of target mRNA (**Figure 6A**). Although small interfering RNAs (siRNAs) are widely used for this purpose, their duplex structure renders them relatively inert to 2′-OH acylation,^30, 58^ and separate strand modification would likely compromise duplex reformation and functional recovery in cells. We therefore selected short hairpin RNA (shRNA) as a more suitable scaffold. shRNAs are single-stranded transcripts containing complementary stem regions connected by a conserved loop sequence recognized by Dicer. Upon processing, Dicer removes the loop to generate a duplex that is incorporated into RISC, where the Argonaute (Ago) protein mediates target mRNA cleavage^59–61^. We hypothesized that **PIC**-mediated acylation of shRNA would disrupt Dicer recognition and processing of the hairpin structure, thereby abolishing downstream gene silencing until reductive decaging restored functionality (**Figure 6A**).

**FIGURE 6.**
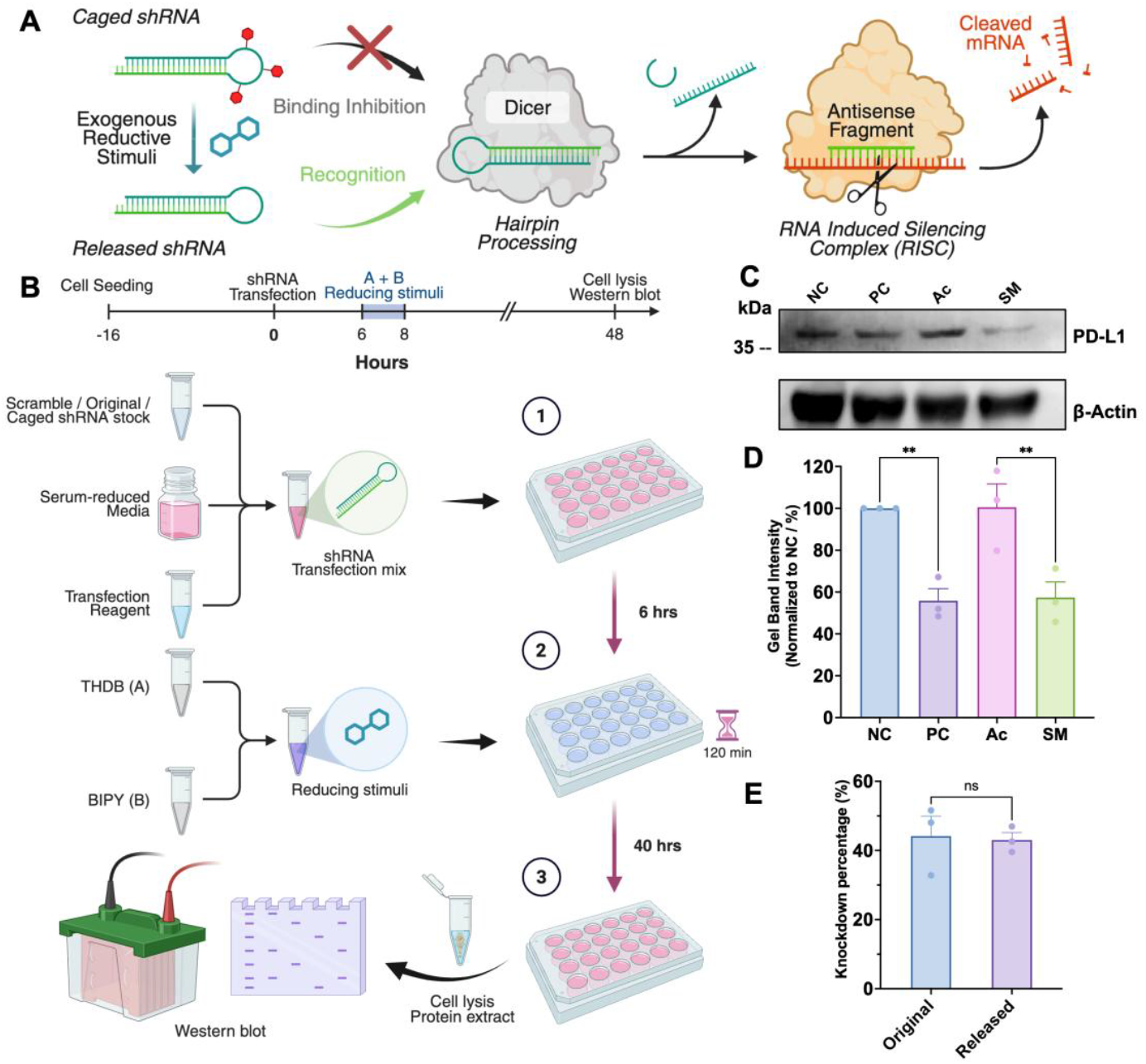
Short hairpin RNA (shRNA) caging strategy for RNAi application. A) Mechanism of shRNA-mediated target gene silencing after reducing stimuli induced uncaging of **PIC** protection. B) Schematic illustration of RNAi experimental design. SNU449 cells were transfected with scramble shRNA, unmodified (mock-treated) or 50 mM **PIC** modified shPD-L1 for 6 h and treated with blank treatment (1X PBS) or mixture of 4 mM A and 1 mM B in 1X PBS for 120 min, then further cultured for 40 h before lysed at 4 °C with RIPA buffer (protease inhibitor added) and analyzed with Western blotting. C) Fluorescent imaging of immunoblot for shPD-L1 uncaging. Cells were transfected with scramble shRNA (NC), unmodified (PC) or 50 mM **PIC** acylated shPD-L1 (Ac, SM) and treated with blank treatment (NC, PC, Ac) or reducing small molecule stimuli (SM). Data are representatives of three replicate experiments. E) Comparison of knockdown efficacy between original shRNA and cage-released shRNA. Knockdown percentages are calculated based on relative intensity difference for each replicate, e.g., 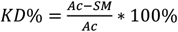for released shRNA. Data are presented as the mean ± SEM of n = 3 independent experi-ments.

Cancer remains one of the most lethal diseases worldwide, in part due to the ability of malignant cells to evade immune surveillance.^62^ A central mechanism of immune evasion involves overexpression of ligands that engage T cell immune check-points, such as programmed death-ligand 1 (PD-L1) on tumor cells binding to programmed cell death protein 1 (PD-1) on T cells, thereby suppressing antitumor immune responses.^63, 64^ RNA-based strategies for modulating immune checkpoint expression have emerged as a promising approach to restore immune activity.^10, 65^ To evaluate the applicability of our chemically controllable RNA platform, we designed a shRNA targeting a sequence within the PD-L1 open reading frame, incorporating a CUCGAG loop for Dicer recognition (**Figure S10**). Functional validation was performed in SNU449 cells, a grade II–III/IV hepatocellular carcinoma line with confirmed PD-L1 expression,^66^ establishing a model system to assess the potential of reversible RNA acylation as a chemical caging strategy for precise, stimulus-responsive regulation of gene expression in living cells.

Cell-based experiments were conducted over a 48 h timeline (**Figure 6B**, upper). shRNA targeting PD-L1 (shPD-L1) was modified with 50 mM **PIC** and purified to afford a quantitatively recovered caged construct with confirmed integrity by gel electrophoresis (**Figure S11**). SNU449 hepatocellular carcinoma cells were seeded overnight and transfected with 1 μg of **PIC**-caged shPD-L1. Scrambled shRNA served as a negative control, and unmodified shPD-L1 as a positive control. Six hours post-transfection, cells were treated with 4 mM **A** and 1 mM **B** for 120 min, washed, and further cultured in RPMI 1640 medium supplemented with FBS for 40 h. Total cellular protein was then extracted and analyzed by Western blotting following separation on 12% SDS-PAGE gels (**Figure 6B**, lower). Immunoblot analysis (**Figure 6C**) revealed a marked decrease in PD-L1 expression upon treatment with the **A**/**B** reducing pair. Densitometric quantification (**Figure 6D** and **S12**) demonstrated significant knockdown in the decaged group relative to both the caged and scrambled controls. In contrast, minimal knockdown was observed in the presence of the caging modification. Comparison of relative knockdown efficiencies indicated no statistically significant difference between the decaged shRNA and the unmodified positive control across triplicates, confirming effective restoration of gene-silencing activity (**Figure 6E**). These findings establish **PIC**-mediated reversible acylation as a controllable RISC-inducing strategy, enabling target mRNA cleavage only upon exposure to the THDB/BIPY reducing system and thereby providing a chemically gated on– off switch for gene silencing. Collectively, these results highlight the potential of this reversible acylation platform as a chemically controllable caging toolkit for nucleic acid–based therapeutic strategies, enabling temporal and stimulus-specific regulation of RNA-mediated gene silencing in living cells.

## CONCLUSIONS

We report a chemically triggered, control–release strategy for RNA based on 2′-OH modification with a nitro-containing carbamate. Upon treatment with the THDB–BIPY reducing pair at low millimolar concentrations, the modification is efficiently and rapidly removed within minutes. Installation of this group sterically disrupts RNA folding and intermolecular interactions, thereby suppressing native function, while reductive cleavage restores structure and activity. This approach is broadly applicable across diverse RNA classes, including fluorogenic aptamers, single-guide RNAs, short hairpin RNAs, and long mRNAs. In *cellulo* studies confirmed biocompatibility and selective functional recovery, while *in vitro* assays showed bioorthogonality, with minimal reactivity toward endogenous reductants or commonly used reducing agents. Collectively, this platform enables stimulus-responsive, exogenously controlled regulation of RNA function, providing an orthogonal toolkit for reversible RNA modulation that complements existing strategies based on endogenous-responsive protection, with potential applications in research and therapeutic development.

## ASSOCIATED CONTENT

### Supporting Information

The Supporting Information is available free of charge on the ACS Publications website.

Materials and reagents used, general procedures and methods of chemical synthesis, in vitro validation and cell experiments, supplementary Figures, NMR and mass spectrometry spectra (docx).

## ACKNOWLEDGMENT

Figures were created with BioRender. The author thanks Zhengyu Lin for advice in Western blotting experiments. HeLa cell line was a generous gift from Prof. Shao Q. Yao and Fengfei Miao, National University of Singapore. SNU449 cell line was a generous gift from Prof. Ang Wee Han and Hanqing Pang, National University of Singapore.

## ABBREVIATIONS

nt: (number of) nucleotides
NAD(P)H: nicotinamide adenine di-nucleotide (phosphate) reduced form
GSH: glutathione
DMEM: Dulbecco’s Modified Eagle Medium
RPMI 1640: Roswell Park Memorial Institute 1640
FBS: fetal bovine serum
ANOVA: analysis of variance
DMSO: dimethyl sulfoxide
RNAi: RNA interference
NC: negative control
PC: positive control

## REFERENCES

(1) Das, S.; Vera, M.; Gandin, V.; Singer, R. H.; Tutucci, E. Intra-cellular mRNA transport and localized translation. Nat Rev Mol Cell Biol 2021, 22 (7), 483–504. DOI: 10.1038/s41580-021-00356-8.

(2) Jia, X.; He, X.; Huang, C.; Li, J.; Dong, Z.; Liu, K. Protein translation: biological processes and therapeutic strategies for human diseases. Signal Transduct Target Ther 2024, 9 (1), 44. DOI: 10.1038/s41392-024-01749-9.

(3) Anastasiadou, E.; Jacob, L. S.; Slack, F. J. Non-coding RNA networks in cancer. Nat Rev Cancer 2018, 18 (1), 5–18. DOI: 10.1038/nrc.2017.99.

(4) Holoch, D.; Moazed, D. RNA-mediated epigenetic regulation of gene expression. Nat Rev Genet 2015, 16 (2), 71–84. DOI: 10.1038/nrg3863.

(5) Chen, L. L.; Kim, V. N. Small and long non-coding RNAs: Past, present, and future. Cell 2024, 187 (23), 6451–6485. DOI: 10.1016/j.cell.2024.10.024.

(6) Zou, Z.; Wei, J.; He, C. New horizons of regulatory RNA. Fundam Res 2023, 3 (5), 760–762. DOI: 10.1016/j.fmre.2022.12.001.

(7) Wang, Y.-S.; Kumari, M.; Chen, G.-H.; Hong, M.-H.; Yuan, J. P.-Y.; Tsai, J.-L.; Wu, H.-C. mRNA-based vaccines and therapeutics: an in-depth survey of current and upcoming clinical applications. Journal of Biomedical Science 2023, 30 (1). DOI: 10.1186/s12929-023-00977-5.

(8) Zhu, Y.; Zhu, L.; Wang, X.; Jin, H. RNA-based therapeutics: an overview and prospectus. Cell Death & Disease 2022, 13 (7). DOI: 10.1038/s41419-022-05075-2.

(9) Collotta, D.; Bertocchi, I.; Chiapello, E.; Collino, M. Antisense oligonucleotides: a novel Frontier in pharmacological strategy. Frontiers in Pharmacology 2023, 14. DOI: 10.3389/fphar.2023.1304342.

(10) Chehelgerdi, M.; Chehelgerdi, M. The use of RNA-based treatments in the field of cancer immunotherapy. Molecular Cancer 2023, 22 (1). DOI: 10.1186/s12943-023-01807-w.

(11) Saw, P. E.; Song, E. Advancements in clinical RNA therapeutics: Present developments and prospective outlooks. Cell Rep Med 2024, 5 (5), 101555. DOI: 10.1016/j.xcrm.2024.101555.

(12) Kariko, K.; Buckstein, M.; Ni, H.; Weissman, D. Suppression of RNA recognition by Toll-like receptors: the impact of nucleoside modification and the evolutionary origin of RNA. Immunity 2005, 23 (2), 165–175. DOI: 10.1016/j.immuni.2005.06.008.

(13) Kariko, K.; Muramatsu, H.; Welsh, F. A.; Ludwig, J.; Kato, H.; Akira, S.; Weissman, D. Incorporation of pseudouridine into mRNA yields superior nonimmunogenic vector with increased translational capacity and biological stability. Mol Ther 2008, 16 (11), 1833–1840. DOI: 10.1038/mt.2008.200.

(14) Li, Y.; Yi, Y.; Gao, X.; Wang, X.; Zhao, D.; Wang, R.; Zhang, L. S.; Gao, B.; Zhang, Y.; Zhang, L.; et al. 2’-O-methylation at internal sites on mRNA promotes mRNA stability. Mol Cell 2024, 84 (12), 2320–2336 e2326. DOI: 10.1016/j.molcel.2024.04.011.

(15) Kuhn, A. N.; Diken, M.; Kreiter, S.; Selmi, A.; Kowalska, J.; Jemielity, J.; Darzynkiewicz, E.; Huber, C.; Tureci, O.; Sahin, U. Phosphorothioate cap analogs increase stability and translational efficiency of RNA vaccines in immature dendritic cells and induce superior immune responses in vivo. Gene Ther 2010, 17 (8), 961–971. DOI: 10.1038/gt.2010.52.

(16) Strzelecka, D.; Smietanski, M.; Sikorski, P. J.; Warminski, M.; Kowalska, J.; Jemielity, J. Phosphodiester modifications in mRNA poly(A) tail prevent deadenylation without compromising protein expression. RNA 2020, 26 (12), 1815–1837. DOI: 10.1261/rna.077099.120.

(17) Geri, J. B.; Oakley, J. V.; Reyes-Robles, T.; Wang, T.; McCarver, S. J.; White, C. H.; Rodriguez-Rivera, F. P.; Parker, D. L., Jr.; Hett, E. C.; Fadeyi, O. O.; et al. Microenvironment mapping via Dexter energy transfer on immune cells. Science 2020, 367 (6482), 1091–1097. DOI: 10.1126/science.aay4106.

(18) Kovachka, S.; Wang, J.; Taghavi, A.; Jia, Y.; Asaba, T.; Wolff, K. C.; Martin, M.; Yang, X.; Meyer, S. M.; Ottilie, S.; et al. Covalent Probes Reveal Small-Molecule Binding Pockets in Structured RNA and Enable Bioactive Compound Design. J Am Chem Soc 2025, 147 (41), 37460–37479. DOI: 10.1021/jacs.5c11898.

(19) Branon, T. C.; Bosch, J. A.; Sanchez, A. D.; Udeshi, N. D.; Svinkina, T.; Carr, S. A.; Feldman, J. L.; Perrimon, N.; Ting, A. Y. Efficient proximity labeling in living cells and organisms with TurboID. Nat Biotechnol 2018, 36 (9), 880–887. DOI: 10.1038/nbt.4201.

(20) Lei, Y.; Zheng, M.; Chen, P.; Seng Ng, C.; Peng Loh, T.; Liu, H. Linker Design for the Antibody Drug Conjugates: A Comprehensive Review. ChemMedChem 2025, 20 (15), e202500262. DOI: 10.1002/cmdc.202500262.

(21) Fralish, Z.; Chen, A.; Khan, S.; Zhou, P.; Reker, D. The landscape of small-molecule prodrugs. Nat Rev Drug Discov 2024, 23 (5), 365–380. DOI: 10.1038/s41573-024-00914-7.

(22) Su, Z.; Xiao, D.; Xie, F.; Liu, L.; Wang, Y.; Fan, S.; Zhou, X.; Li, S. Antibody-drug conjugates: Recent advances in linker chemistry. Acta Pharm Sin B 2021, 11 (12), 3889–3907. DOI: 10.1016/j.apsb.2021.03.042.

(23) Do, T. C.; Lau, J. W.; Sun, C.; Liu, S.; Kha, K. T.; Lim, S. T.; Oon, Y. Y.; Kwan, Y. P.; Ma, J. J.; Mu, Y.; et al. Hypoxia deactivates epigenetic feedbacks via enzyme-derived clicking proteolysis-targeting chimeras. Sci Adv 2022, 8 (50), eabq2216. DOI: 10.1126/sciadv.abq2216.

(24) Wu, C. S.; Cheng, L. Recent Advances towards the Reversible Chemical Modification of Proteins. Chembiochem 2023, 24 (2), e202200468. DOI: 10.1002/cbic.202200468.

(25) Chen, Y.; Dai, C.; Han, J.; Xing, Y.; Yin, F.; Li, Z. Recent Chemical Biology Insights Towards Reversible Stapled Peptides. Chembiochem 2025, 26 (9), e202500052. DOI: 10.1002/cbic.202500052.

(26) Tavakoli, A.; Min, J. H. Photochemical modifications for DNA/RNA oligonucleotides. RSC Adv 2022, 12 (11), 6484–6507. DOI: 10.1039/d1ra05951c.

(27) Ducho, C. Enzymatically Cleavable siRNA Prodrugs: a New Paradigm for the Intracellular Delivery of RNA-Based Therapeutics. ChemMedChem 2015, 10 (10), 1625–1627. DOI: 10.1002/cmdc.201500279.

(28) Wang, S.; Saneyoshi, H.; Xu, P.; Oguri, N.; Yamashita, A.; Xu, Y. Manipulating DNA and RNA structures via click-to-release caged nucleic acids for biological and biomedical applications. Nucleic Acids Res 2025, 53 (12). DOI: 10.1093/nar/gkaf571.

(29) Obexer, R.; Nassir, M.; Moody, E. R.; Baran, P. S.; Lovelock, S. L. Modern approaches to therapeutic oligonucleotide manufacturing. Science 2024, 384 (6692), eadl4015. DOI: 10.1126/science.adl4015.

(30) Velema, W. A.; Kool, E. T. The chemistry and applications of RNA 2′-OH acylation. Nature Reviews Chemistry 2019, 4 (1), 22–37. DOI: 10.1038/s41570-019-0147-6.

(31) Fang, L.; Xiao, L.; Jun, Y. W.; Onishi, Y.; Kool, E. T. Reversible 2′-OH acylation enhances RNA stability. Nature Chemistry 2023, 15 (9), 1296–1305. DOI: 10.1038/s41557-023-01246-6.

(32) Jash, B.; Kool, E. T. Conjugation of RNA via 2’-OH acylation: Mechanisms determining nucleotide reactivity. Chem Commun (Camb) 2022, 58 (22), 3693–3696. DOI: 10.1039/d2cc00660j.

(33) Smola, M. J.; Rice, G. M.; Busan, S.; Siegfried, N. A.; Weeks, K. M. Selective 2’-hydroxyl acylation analyzed by primer extension and mutational profiling (SHAPE-MaP) for direct, versatile and accurate RNA structure analysis. Nat Protoc 2015, 10 (11), 1643–1669. DOI: 10.1038/nprot.2015.103.

(34) Flynn, R. A.; Zhang, Q. C.; Spitale, R. C.; Lee, B.; Mumbach, M. R.; Chang, H. Y. Transcriptome-wide interrogation of RNA secondary structure in living cells with icSHAPE. Nat Protoc 2016, 11 (2), 273–290. DOI: 10.1038/nprot.2016.011.

(35) Spitale, R. C.; Crisalli, P.; Flynn, R. A.; Torre, E. A.; Kool, E. T.; Chang, H. Y. RNA SHAPE analysis in living cells. Nat Chem Biol 2013, 9 (1), 18–20. DOI: 10.1038/nchembio.1131.

(36) Kha, T. K.; Zhao, Y.; Zhu, R. Y. Site-Selective Modification and Labeling of Native RNA. Chemistry 2025, 31 (12), e202404244. DOI: 10.1002/chem.202404244.

(37) Kha, T. K.; Zhang, T.; Kunchur, N.; Zhao, Y.; Guo, J.; Chen, S.; Zhu, R. Y. Site-Selective RNA Modification by Programmable DNA-Small Molecule Conjugates. Angew Chem Int Ed Engl 2025, 64 (52), e15411. DOI: 10.1002/anie.202515411.

(38) Pratihar, S.; Zhong, W.; Feng, S.; Chatterjee, S.; Kool, E. T. Sequence-Specific Installation of Aryl Groups in RNA via DNA-Catalyst Conjugates. Angew Chem Int Ed Engl 2025, 64 (46), e202515681. DOI: 10.1002/anie.202515681.

(39) Kadina, A.; Kietrys, A. M.; Kool, E. T. RNA Cloaking by Reversible Acylation. Angew Chem Int Ed Engl 2018, 57 (12), 3059–3063. DOI: 10.1002/anie.201708696.

(40) Velema, W. A.; Kietrys, A. M.; Kool, E. T. RNA Control by Photoreversible Acylation. J Am Chem Soc 2018, 140 (10), 3491–3495. DOI: 10.1021/jacs.7b12408.

(41) Guo, J.; Chen, S.; Onishi, Y.; Shi, Q.; Song, Y.; Mei, H.; Chen, L.; Kool, E. T.; Zhu, R. Y. RNA Control via Redox-Responsive Acylation. Angew Chem Int Ed Engl 2024, 63 (21), e202402178. DOI: 10.1002/anie.202402178.

(42) Guo, J.; Zhang, S.; Kha, T. K.; Hu, C.; Zhu, R. Y. Accelerating Responsive RNA Release Through Structural Optimization of Disulfide-Containing Acyl Groups. Angew Chem Int Ed Engl 2025, 64 (35), e202507581. DOI: 10.1002/anie.202507581.

(43) Denny, W. A. Nitroaromatic Hypoxia-Activated Prodrugs for Cancer Therapy. Pharmaceuticals (Basel) 2022, 15 (2). DOI: 10.3390/ph15020187.

(44) Weng, C.; Yang, H.; Loh, B. S.; Wong, M. W.; Ang, W. H. Targeting Pathogenic Formate-Dependent Bacteria with a Bioinspired Metallo-Nitroreductase Complex. J Am Chem Soc 2023, 145 (11), 6453–6461. DOI: 10.1021/jacs.3c00237.

(45) Parkinson, G. N.; Skelly, J. V.; Neidle, S. Crystal structure of FMN-dependent nitroreductase from Escherichia coli B: a prodrug-activating enzyme. J Med Chem 2000, 43 (20), 3624–3631. DOI: 10.1021/jm000159m.

(46) Hacıoğlu, N.; Güngör, T.; Tokay, E.; Gülhan, Ü.G.; Çelik, A.; Ay, M.; Köçkar, F. Prodrugs for Nitroreductase Based Cancer Therapy-5: Development of Trinitroaniline Prodrugs/Ssap-NtrB Combinations for Liver Cancer Using Intracellular and Extracellular Conditions. ChemistrySelect 2021, 6 (25), 6315–6323. DOI: 10.1002/slct.202101115.

(47) Wilson, W. R.; Hay, M. P. Targeting hypoxia in cancer therapy. Nature Reviews Cancer 2011, 11 (6), 393–410. DOI: 10.1038/nrc3064.

(48) Mitchell, D. J.; Minchin, R. F. E. coli nitroreductase/CB1954 gene-directed enzyme prodrug therapy: role of arylamine N-acet-lytransferase 2. Cancer Gene Ther 2008, 15 (11), 758–764. DOI: 10.1038/cgt.2008.47.

(49) Qi, J. Q.; Jiao, L. DFT Study on the Mechanism of 4,4’-Bipyridine-Catalyzed Nitrobenzene Reduction by Diboron(4) Compounds. J Org Chem 2020, 85 (21), 13877–13885. DOI: 10.1021/acs.joc.0c01963.

(50) Hosoya, H.; Misal Castro, L. C.; Sultan, I.; Nakajima, Y.; Ohmura, T.; Sato, K.; Tsurugi, H.; Suginome, M.; Mashima, K. 4,4′-Bipyridyl-Catalyzed Reduction of Nitroarenes by Bis(neopentylglycolato)diboron. Organic Letters 2019, 21 (24), 9812–9817. DOI: 10.1021/acs.orglett.9b03419.

(51) Wang, Q.; Song, Y.; Yuan, S.; Zhu, Y.; Wang, W.; Chu, L. Prodrug activation by 4,4’-bipyridine-mediated aromatic nitro reduction. Nat Commun 2024, 15 (1), 8643. DOI: 10.1038/s41467-024-52604-y.

(52) Velema, W. A.; Kool, E. T. Water-Soluble Leaving Group Enables Hydrophobic Functionalization of RNA. Org Lett 2018, 20 (20), 6587–6590. DOI: 10.1021/acs.orglett.8b02938.

(53) Chen, X.; Zhang, D.; Su, N.; Bao, B.; Xie, X.; Zuo, F.; Yang, L.; Wang, H.; Jiang, L.; Lin, Q.; et al. Visualizing RNA dynamics in live cells with bright and stable fluorescent RNAs. Nat Biotechnol 2019, 37 (11), 1287–1293. DOI: 10.1038/s41587-019-0249-1.

(54) Huang, K.; Chen, X.; Li, C.; Song, Q.; Li, H.; Zhu, L.; Yang, Y.; Ren, A. Structure-based investigation of fluorogenic Pepper aptamer. Nat Chem Biol 2021, 17 (12), 1289–1295. DOI: 10.1038/s41589-021-00884-6.

(55) Ran, F. A.; Hsu, P. D.; Wright, J.; Agarwala, V.; Scott, D. A.; Zhang, F.; Ran, F. A.; Hsu, P. D.; Wright, J.; Agarwala, V.; et al. Genome engineering using the CRISPR-Cas9 system. Nature Protocols 2013 8:11 2013-10-24, 8 (11). DOI: 10.1038/nprot.2013.143.

(56) Li, T.; Yang, Y.; Qi, H.; Cui, W.; Zhang, L.; Fu, X.; He, X.; Liu, M.; Li, P. F.; Yu, T. CRISPR/Cas9 therapeutics: progress and prospects. Signal Transduct Target Ther 2023, 8 (1), 36. DOI: 10.1038/s41392-023-01309-7.

(57) Brito Querido, J.; Diaz-Lopez, I.; Ramakrishnan, V. The molecular basis of translation initiation and its regulation in eukaryotes. Nat Rev Mol Cell Biol 2024, 25 (3), 168–186. DOI: 10.1038/s41580-023-00624-9.

(58) Habibian, M.; Velema, W. A.; Kietrys, A. M.; Onishi, Y.; Kool, E. T. Polyacetate and Polycarbonate RNA: Acylating Reagents and Properties. Org Lett 2019, 21 (14), 5413–5416. DOI: 10.1021/acs.orglett.9b01526.

(59) Lambeth, L. S.; Smith, C. A. Short hairpin RNA-mediated gene silencing. Methods Mol Biol 2013, 942, 205–232. DOI: 10.1007/978-1-62703-119-6_12.

(60) McIntyre, G. J.; Fanning, G. C. Design and cloning strategies for constructing shRNA expression vectors. BMC Biotechnol 2006, 6, 1. DOI: 10.1186/1472-6750-6-1.

(61) McIntyre, G. J.; Yu, Y. H.; Lomas, M.; Fanning, G. C. The effects of stem length and core placement on shRNA activity. BMC Mol Biol 2011, 12, 34. DOI: 10.1186/1471-2199-12-34.

(62) Global Burden of Disease Cancer, C.; Kocarnik, J. M.; Compton, K.; Dean, F. E.; Fu, W.; Gaw, B. L.; Harvey, J. D.; Henrikson, H. J.; Lu, D.; Pennini, A.; et al. Cancer Incidence, Mortality, Years of Life Lost, Years Lived With Disability, and Disability-Adjusted Life Years for 29 Cancer Groups From 2010 to 2019: A Systematic Analysis for the Global Burden of Disease Study 2019. JAMA Oncol 2022, 8 (3), 420–444. DOI: 10.1001/jamaoncol.2021.6987.

(63) Tufail, M.; Jiang, C. H.; Li, N. Immune evasion in cancer: mechanisms and cutting-edge therapeutic approaches. Signal Transduct Target Ther 2025, 10 (1), 227. DOI: 10.1038/s41392-025-02280-1.

(64) He, X.; Xu, C. Immune checkpoint signaling and cancer immunotherapy. Cell Res 2020, 30 (8), 660–669. DOI: 10.1038/s41422-020-0343-4.

(65) Choi, Y.; Seok, S. H.; Yoon, H. Y.; Ryu, J. H.; Kwon, I. C. Advancing cancer immunotherapy through siRNA-based gene silencing for immune checkpoint blockade. Adv Drug Deliv Rev 2024, 209, 115306. DOI: 10.1016/j.addr.2024.115306.

(66) Wu, M.; Xia, X.; Hu, J.; Fowlkes, N. W.; Li, S. WSX1 act as a tumor suppressor in hepatocellular carcinoma by downregulating neoplastic PD-L1 expression. Nat Commun 2021, 12 (1), 3500. DOI: 10.1038/s41467-021-23864-9.

