## Supporting Information for "Nitro Reduction-Based RNA Control and Ultrafast Release"

### Table of contents

**Materials and Reagents**

**General Procedures and Methods**

- Chemical Synthesis of Acylating Agents
- Methods for *in vitro* Validation
- Methods for Cell Experiments

**Supplementary Figures**

**NMR Spectroscopy and Mass Spectrometry**

### Materials

Chemical and biochemical reagents are purchased from commercially available sources (S. Table 1) and was used without further purification.

**Supplementary Table 1. List of Chemical Reagents used in this work.**

| Reagents | Source | Catalog Number |
| --- | --- | --- |
| Imidazole-2-carboxylic acid | BLDpharm | BD41338 |
| N,N,N′-Trimethylethylenediamine | BLDpharm | BD141803 |
| 1,1’-carbonyldiimidazole | Sigma-Aldrich | #115533 |
| 1H-imidazole | BLDpharm | BD40902 |
| 2-chloro-1H-imidazole | BLDpharm | BD2066 |
| 4-nitrobenzyl chloroformate | Sigma-Aldrich | #222801 |
| Tetrahydroxydiboron **(THDB, “A”)** | Sigma-Aldrich | #754242 |
| 4,4’-bipyridine **(BIPY, “B”)** | Sigma-Aldrich | #289426 |
| HEPES | Sigma-Aldrich | #H3375 |
| MOPS | Sigma-Aldrich | #M1254 |
| 3-hydroxypicolinic acid | Sigma-Aldrich | #56197 |
| Ammonium citrate dibasic | Sigma-Aldrich | #09833 |
| Magnesium chloride | BLDpharm | BD136984 |
| Ethanol, absolute | VWR Chemicals | VWRC20821.321 |
| Urea | Sigma-Aldrich | #U1250 |
| Ammonium Persulfate | Bio-Rad | #1610700 |
| SureCast™ TEMED | Thermo Fisher Scientific | HC2006 |
| SureCast™ Acrylamide Solution (40%) | Thermo Fisher Scientific | HC2040 |
| EDTA tetrasodium | BLDpharm | BD105667 |
| Ammonium acetate | Sigma-Aldrich | #238074 |

Short DNA or RNA oligonucleotides are directly purchased from Integrated DNA Technologies without further purification or synthesized via *in vitro* transcription (IVT). shRNA was designed from siRNA-targeting regions, analyzed with GPP Web Portal (Broad Institute)[1] and purchased from Genscript Biotech.

**Supplementary Table 2. DNA/RNA sequences.**

| Name | Sequence |
|---|---|
| 18-mer FAM-RNA | /6-FAM/AUC CUG CCG ACU ACG CCA |
| 39-mer RNA | UAU UUA GUC CCA GGA UCA CGU UAA CGU ACU ACA UCA CUA |
| Full complement 39-mer DNA | TAG TGA TGT AGT ACG TTA ACG TGA TCC TGG GAC TAA ATA |
| T7 promoter | TAA TAC GAC TCA CTA TAG GG |
| shPD-L1 | CCG GCC UAC UGG CAU UUG CUG AAC GCU CGA GCG UUC AGC AAA UGC CAG UAG GUU UUU G |
| Scramble shPD-L1 | CCG GCA CUU GUC GAC CGU AUG CUA CCU CGA GGU AGC AUA CGG UCG ACA AGU GUU UUU G |
| F30-pepper RNA (IVT) | GGG UUG CCA UGU GUA UGU GGG UUC GCC CAC AUA CUC UGA UGA UCC CCA AUC GUG GCG UGU CGG CCU GCU UCG GCA GGC ACU GGC GCC GGG AUC AUU CAU GGC AA |
| 131-mer guide RNA (IVT) | GUC CUC AGA UUU AGU AUU CAG AGU UGU AGC UCC CUU UCU CGA AAG AGA ACC GUU GCU ACA AUA AGG CCG UCU GAA AAG AUG UGC CGC AAC GCU CUG CCC CUU AAA GCU UCU GCU UUA AGG GGC AUC GUU UA |

**Supplementary Table 3. Sequence information of proteins.**

| Name | Sequence |
|---|---|
| NmeCas9 | MAAFKPNSINYILGLDIGIASVGWAMVEIDEEENPIRLIDLGVRVFERAE<br>VPKTGDSLAMARRLARSVRRLTRRRAHRLLRTRRLLKREGVLQAANF<br>DENGLIKSLPNTPWQLRAAALDRKLTPLEWSAVLLHLIKHRGYLSQRKN<br>EGETADKELGALLKGVAGNAHALQTGDFRTPAELALNKFEKESGHIRN<br>QRSDYSHTFSRKDLQAELILLFEKQKEFGNPHVSGGLKEGIETLLMTQ<br>RPALSGDAVQKMLGHCTFEPAEPKAAKNTYTAERFIWLTKLNNLRILEQ<br>GSERPLTDTERATLMDEPYRKSKLTYAQARKLLGLEDTAFFKGLRYGK<br>DNAEASTLMEMKAYHAISRALEKEGLKDKKSPLNLSPELQDEIGTAFSL<br>FKTDEDITGRLKDRIQPEILEALLKHISFDKFVQISLKALRRIVPLMEQGK<br>RYDEACAEIYGDHYGKKNTEEKIYLPPIPADEIRNPVVLRALSQARKVIN<br>GVVRRYGSPARIHIETAREVGKSFKDRKEIEKRQEENRKDREKAAAKF<br>REYFPNFVGEPKSKDILKLRLYEQQHGKCLYSGKEINLGRLNEKGYVEI<br>DHALPFSRTWDDSFNNKVLVLGSENQNKGNQTPYEYFNGKDNSREW<br>QEFKARVETSRFPRSKKQRILLQKFDEDGFKERNLNDTRYVNRFLCQF<br>VADRMRLTGKGKKRVFASNGQITNLLRGFWGLRKVRAENDRHHALDA<br>VVVACSTVAMQQKITRFVRYKEMNAFDGKTIDKETGEVLHQKTHFPQP<br>WEFFAQEVMIRVFGKPDGKPEFEEADTLEKLRTLLAEKLSSRPEAVHE<br>YVTPLFVSRAPNRKMSGQGHMETVKSAKRLDEGVSVLRVPLTQLKLK<br>DLEKMVNREREPKLYEALKARLEAHKDDPAKAFAEPFYKYDKAGNRT<br>QQVKAVRVEQVQKTGVWVRNHNGIADNATMVRVDVFEKGDKYYLVPI<br>YSWQVAKGILPDRAVVQGKDEEDWQLIDDSFNFKFSLHPNDLVEVITK<br>KARMFGYFASCHRGTGNINIRIHDLDHKIGKNGILEGIGVKTALSFQKYQ<br>IDELGKEIRPCRLKKRPPVR |
| EGFP | MVSKGEELFTGVVPILVELDGDVNGHKFSVSGEGEGDATYGKLTLKFIC<br>TTGKLPVPWPTLVTTLTYGVQCFSRYPDHMKQHDFFKSAMPEGYVQE<br>RTIFFKDDGNYKTRAEVKFEGDTLVNRIELKGIDFKEDGNILGHKLEYNY<br>NSHNVYIMADKQKNGIKVNFKIRHNIEDGSVQLADHYQQNTPIGDGPV<br>LLPDNHYLSTQSALSKDPNEKRDHMVLLEFVTAAGITLGMDELYK |

##### Molecular and Cell Biology Materials and Kits

384 Well Black Plate, 96 Well Plate, 24 Well Plate, 12 Well Plate, 6 Well Plate, Dulbecco's Modified Eagle's Medium/High Glucose (phenol red), Opti-MEM™ I Reduced Serum Medium (no phenol red), 0.25% Trypsin-EDTA (phenol red), VALUE FBS (Fetal Bovine Sera), Penicillin-Streptomycin (10,000 U/mL), Bis-benzimide H33342 trihydrochloride (Hoechst 33342), RNaseOUT™ Recombinant Ribonuclease Inhibitor, RiboRuler High Range RNA Ladder, Lipofectamine™ 3000 Transfection Reagent and Lipofectamine™ MessengerMAX™ Transfection Reagent, Pierce™ BCA Protein Assay Kit, RIPA Lysis Buffer, Pierce Protease Inhibitor Mini Tablets, PVDF Transfer Membranes 0.45 µm were purchased from Thermo Fisher Scientific. CAG EGFP mRNA (N1-Me-Pseudo UTP) was purchased from Vazyme. HyPure molecular biology-grade water and HyClone™ RPMI 1640 liquid media was purchased from HyClone. 10X Tris-Acetate-EDTA (TBE) pH 8.0, 10X Tris-Borate-EDTA (TBE) pH 7.4, 10X Phosphate Buffered Saline (PBS) pH 7.4, 20X Tris Buffered Saline (TBS), 0.5M Tris Buffer, pH 6.8, 1.0M Tris Buffer, pH 8.0, 3 M sodium acetate (NaOAc) pH 5.2, 10X Tris Glycine (TG) Buffer, 10X Tris Glycine-Sodium Dodecyl Sulfate (TG-SDS) Buffer, and 6x DNA Loading Dye were purchased from Axil Scientific. HiScribe® T7 High Yield RNA Synthesis Kit was purchased from New England Biolabs. RNA Clean & Concentrator™ was purchased from Zymo Research. Wheat Germ Extract, 1.0 M KOAc and Complete Amino Acid Mixture were purchased from Promega.

##### Instrumentation

Purification of imidazole-2-amide through reverse phase was conducted on CombiFlash NextGen 100 (Teledyne LABS), and lyophilization of aqueous product was performed using Labconco FreeZone Benchtop Freeze Dryer (-50 °C, 2.5 L). $^{1}$H NMR and $^{13}$C NMR spectra were collected on Bruker AMX400 (400 MHz) and Bruker AMX500 (500 MHz) spectrometer. High-resolution mass spectrometry (HR-MS) was obtained using a Finnigan/MAT 95XL-T spectrometer. Nucleic acid concentrations were measured using NanoDrop Lite Plus Spectrophotometer (Thermo Fisher Scientific). Short RNA oligonucleotides were analyzed by MALDI-TOF mass spectrometer (SpiralTOF JMS-S3000, JEOL). Visualization of gel electrophoresis, western blotting and aptamer fluorescence was conducted either on GelDoc Go Gel Imaging System (Bio-Rad) or Typhoon 9410 fluorescence gel scanner (GE Amersham). Aptamer fluorescence spectra, EGFP signal intensity from *in vitro* translation and BCA-based protein concentration quantification were recorded on Varioskan LUX Multimode Microplate Reader (Thermo Fisher Scientific). Melting point experiment was conducted on Cary 3500 Compact UV-Vis Spectrophotometer (Agilent). Confocal laser scanning microscopy imaging was performed on FLUOVIEW FV3000 (Olympus). Flow Cytometry experiments were carried out on a CytoFLEX LX (Beckman Coulter).

### METHOD

#### Synthesis of acylating agents

Synthesis of N-(2-(dimethylamino)ethyl)-N-methyl-1H-imidazole-2-carboxamide (denote as **imidazole-2-amide**) was conducted following previous report[2].

#### Synthesis of p-nitrobenzyl imidazole carbamate (1)

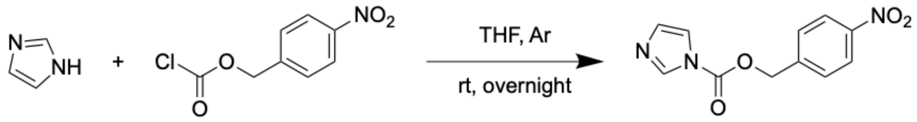


To a stirred solution of 136 mg 1H-imidazole in 4 mL anhydrous THF, 216mg p-nitrobenzyl chloroformate in 0.5 mL anhydrous THF was added dropwise under Ar and stirred overnight. The solid residue was directly filtered, and the filtrate was concentrated under reduced pressure and dried under high vacuum for 2 h. The resulting mixture was dissolved in DMSO-d6 yielding a 2000 uL stock of 500 mM acylating agent and was directly proceeded with acylation experiments without further purification. $^1$H NMR (400 MHz, DMSO) δ 8.35 (t, J = 1.1 Hz, 1H), 8.28 – 8.22 (m, 2H), 7.82 – 7.75 (m, 2H), 7.67 (t, J = 1.5 Hz, 1H), 7.10 (dd, J = 1.7, 0.9 Hz, 1H), 5.59 (s, 2H). $^{13}$C NMR (101 MHz, DMSO) δ 148.15, 147.39, 142.34, 137.42, 130.44, 128.81, 123.63, 117.64, 67.80. HRMS (ESI/Q-TOF) m/z: [M + H]+ Calcd for C11H10N3O4 248.0673; Found 248.0659.

#### Synthesis of p-nitrobenzyl 2-chloroimidazole carbamate (2)

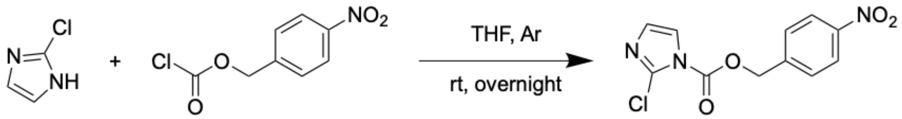


To a stirred solution of 205 mg 2-chloro-1H-imidazole in 4 mL anhydrous THF, 216mg p-nitrobenzyl chloroformate in 0.5 mL anhydrous THF was added dropwise under Ar and stirred overnight. The solid residue was directly filtered, and the filtrate was concentrated under reduced pressure and dried under high vacuum for 2 h. The resulting mixture was dissolved in DMSO-d6 yielding a 2000 uL stock of 500 mM acylating agent and was directly proceeded with acylation experiments. $^{1}H$ NMR (400 MHz, DMSO) δ 8.30 – 8.20 (m, 2H), 7.84 – 7.77 (m, 2H), 7.75 (d, J = 1.9 Hz, 1H), 7.04 (d, J = 2.0 Hz, 1H), 5.58 (s, 2H). $^{13}C$ NMR (101 MHz, DMSO) δ 147.86, 147.56, 142.51, 132.00, 129.37, 128.98, 124.08, 121.84, 68.69. HRMS (ESI/Q-TOF) m/z: [M + H]+ Calcd for C11H9ClN3O4 282.0283; Found 282.0271.

**Synthesis of p-nitrobenzyl imidazole-2-amide carbamate (PIC)**

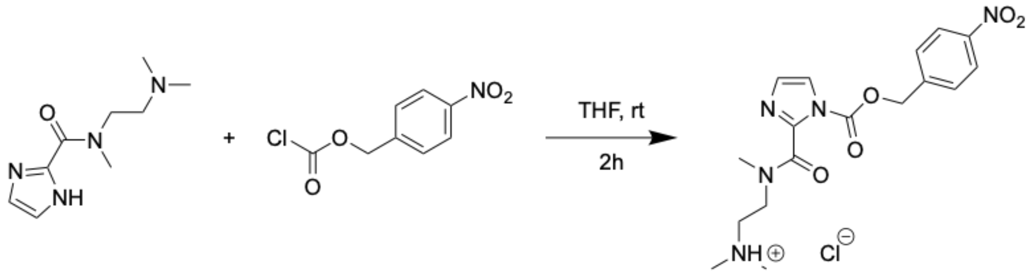

PIC

A solution of 60 / 76 / 102 / 127 mg (0.95, 1.2, 1.6, 2 eq) imidazole-2-amide was dissolved in 1 mL tetrahydrofuran, and 70 mg p-nitrobenzyl chloroformate dissolved in 0.5 mL tetrahydrofuran was added dropwise into the solution. The solution turns turbid immediately after the addition, and the reaction was left to stir at room temperature for 2 hours. Tetrahydrofuran was removed under reduced pressure and dried under high vacuum overnight. Given the high reactivity of product compound, the resulting mixture was dissolved in DMSO-d6 yielding a stock of 500 mM acylating agent (remnants of unreacted I2A not separated) and was directly proceeded with acylation experiments without any purification. $^{1}H$ NMR (400 MHz, DMSO) δ 8.26 (dd, *J* = 8.8, 1.5 Hz, 2H), 7.78 – 7.70 (m, 3H), 7.65 (d, *J* = 8.8 Hz, 1H), 5.55 (d, *J* = 9.0 Hz, 2H), 4.38 (t, *J* = 7.1 Hz, 2H), 3.76 (t, *J* = 6.7 Hz, 1H), 3.40 (t, *J* = 7.1 Hz, 1H), 2.84 (s, 2H), 2.70 (t, *J* = 7.1 Hz, 1H), 2.63 (s, 4H), 2.48 (s, 2H). 13C NMR (101 MHz, DMSO) δ 161.42, 161.25, 159.27, 158.93, 147.55, 147.50, 147.28, 143.04, 142.15, 142.04, 141.95, 141.87, 140.97, 140.71, 129.18, 129.10, 128.72, 127.07, 123.71, 123.65, 123.25, 119.23, 118.83, 68.51, 68.46, 68.35, 61.94, 55.16, 53.80, 53.43, 46.21, 45.65, 43.93, 43.70, 43.45, 43.34, 43.16, 42.77, 36.58, 35.95, 34.37, 32.51, 27.47, 21.80. HRMS (ESI/Q-TOF) m/z: [M + H]+ Calcd for C17H22N5O5 376.1615; Found 376.1616.

**Ethanol precipitation for short RNA oligo purification**

For a 20 µL solution, RNA was precipitated by adding 2 µL (10% V/V) 3 M NaOAc pH 5.2 and 75 µL (375% V/V) ethanol absolute and stored at -80 °C for 16 h. The resulting suspension was centrifuged at 21,100 x g for 60 min at 4 °C, supernatant removed, residue washed twice with 40 µL 75% ethanol and centrifuged each time at 21,100 x g for 12 min. After removing supernatant, residue was resuspended in 10 µL or other desired volume of RNase-free water to yield stock of purified RNA oligo.

For a larger volume of reaction mixture, all amount of reagent use during purification was amplified accordingly (Ex. for 100 µL solution, 10 µL 3 M NaOAc and 375 µL ethanol was added for precipitation and 200 µL 75% ethanol is used for wash).

**General protocol for RNA oligomer acylation**

In a 1.5 mL sterile tube, 200 pmol RNA oligomer was suspended in 16 µL of bio-grade RNase-free water stock solution and equilibrate to 25 °C. 4 µL **PIC** (or **1**, **2**) stock in DMSO (500 mM) was added to the RNA, mixture vortexed briefly to mix well and incubated at 25°C for 120 min. Ethanol precipitation is conducted for purification.

For concentration screening, **PIC** stock was first diluted with anhydrous DMSO to desired stock concentration, then different concentration of **PIC** was added to unmodified RNA oligomer and incubated for 120 min and purified accordingly.

**EGFP mRNA acylation**

In a 1.5 mL sterile tube, 2500 ng EGFP mRNA in MOPS (500 mM, pH 7.5) was reacted with 25 / 30 / 35 mM (*in cellulo* application) / 35 mM (*in vitro* application) **PIC** (final DMSO concentration: 8%) (same amount of DMSO without acylating agent was used for **mock treatment**), mixture pipetted to mix well and incubated for 60 min at room temperature (25 °C) and then isolated by RNA clean and concentrator (Zymo research), RNA concentrated to 15 – 25 µL final volume, sealed and stored at -80 °C.

**General protocol for *in vitro* reduction of modified RNA**

In a 0.2 mL sterile tube, to denoted amount of acylated RNA oligo (typically 500 ng for short oligomers and 300 ng for aptamers) in 1X PBS pH 7.4 (diluted with RNase-free water) was added indicated concentration of **A (tetrahydroxydiboron, THDB)** and **B (4,4'-bipyridine, BIPY)** (Sigma-Aldrich) mixture ([A]:[B] = 4:1) and incubated at 37 °C for different duration and purified either by ethanol precipitation or with RNA clean and concentrator (Zymo research) according to manufacturer's protocol.

***In vitro* transcription synthesis of RNA**

1.5 ug 1:1 mixture of complement DNA and T7 promoter was heated at 95 °C for 5 min and cooled down to room temperature. This mixture was injected as template DNA to HiScribe® T7 High Yield RNA Synthesis Kit (New England Biolabs) and transcribed for 16 h at 37 °C following manufacturer's instructions. Resulting mixture was resolved on an 8% native polyacrylamide gel, extracted with elution buffer (1 mM EDTA, 500 mM ammonium acetate) and isolated with RNA clean and concentrator (Zymo research).

**Melting Point analysis**

In a 0.6 mL sterile tube, 200 pmol 39-mer RNA (original, acylated, reduced) was mixed with 200 pmol fully complementary DNA in 200 µL 1X PBS buffer and transferred to a 300 µL UV-Vis cuvette. 200 µL 1X PBS only was used for blank sample. The melting point program is as follows: Stage I, 25 °C – 90 °C, Ramp rate 40°C/min, 3-min hold at 90 °C; Stage II, 90 °C – 40 °C, Ramp rate 1°C/min. Absorbance measurement at 260 nm was collected from 80 °C to 40 °C with 0.2°C each data point.

**Aptamer Fluorescence**

Based on nanodrop measurement of sample concentration, 50 ng of F30-Pepper RNA aptamer (original, acylated, reduction) was dissolved in 20 µL 1X Folding Buffer (100 mM MOPS pH 7.5, 6 mM $MgCl_2$, 100 mM NaCl) and added 5 µM HBC530 and mixed well. The mixture was incubated in dark for 10 minutes and directly visualized on GE Amersham Typhoon 9410 fluorescence gel scanner. For collection of fluorescence spectra, 20 µL samples were transferred each to a well of black 384-well plate. The emission spectrum was subsequently collected on Varioskan LUX Microplate Reader from 510 to 650 nm, with a fixed excitation wavelength at 485 nm, at 25 ˚C. Intensity of fluorescence was quantified by averaging the readout from 525 nm to 545 nm with reference to negative control (HBC530 in 1X Folding Buffer without aptamer).

**CRISPR-Cas9 assay**

The pRSFDuet target DNA plasmid (36 bp target DNA cloned into the pRSFDuet vector) was linearized by SmaI digestion before the cleavage reaction. First, protected and original gRNAs were incubated with Nme1Cas9 at room temperature for 20 min. Then, the Nme1Cas9 with different modified gRNA complexes were incubated with 100 ng pRSFDuet target DNA and different concentration of A and B (4:1) in 10 μL reaction buffer containing 20 mM HEPES (pH 7.5), 100 mM KCl, 10 mM MgCl2 and 5% glycerol for 20 min at room temperature. Reactions were quenched by adding 2 μl 10x DNA Loading buffer (Vazyme). The reaction products were run on 1% agarose gels stained with Ultra GelRed (Vazyme) for production detection.

***In vitro* translation**

In a 0.2 mL sterile tube, 100 ng of original / modified / released EGFP mRNA (Vazyme) was diluted to 6.4 μL with RNase-free water and incubated at 65 °C for 5 min and flash-cooled on ice. 10 μL Wheat Germ Extract (Promega), 1.6 μL Amino Acid mix (Promega), 0.4 μL RNaseOUT (Thermo Fisher Scientific), and 1.6 μL 1M KOAc (Promega) were then added and mixed well by gentle pipetting. The samples were immediately transferred to a 384-well plate, then incubated in Varioskan LUX Microplate Reader at 25 °C for 120 min. The signal of EGFP was subsequently measured at 530 nm, with a fixed excitation wavelength at 488 nm, at 25 ˚C.

***In cellulo* Validation**

All experiments (unless specified otherwise) were conducted on either HeLa cell line (cervical adenocarcinoma) or SNU449 cell line (hepatocellular carcinoma). HeLa cells were generous gift from Yao S.Q. group, National University of Singapore. SNU449 cells were generous gift from Ang Wee Han group, National University of Singapore.

**Cell culture and seeding**

HeLa cells were grown in T25 cell culture flasks and cultured in DMEM media (+ 10% FBS, + 1% penicillin/streptomycin) in air containing 5% $CO_2$. SNU449 cells were grown in T25 cell culture flasks and cultured in RPMI1640 media (+ 10% FBS) in air containing 5% $CO_2$. Cells were grown until they reached ~ 90% confluency. Cells were harvested by first washing with 3 mL of warm 1x PBS and incubated with 3 mL Trypsin at 37°C for 3 min. After quenching the reaction of trypsin with 6 mL media, mixture was transferred to 15 mL falcon tube and sedimented by centrifugation at 1000 X g for 3 minutes at 25°C. Supernatant was aspirated and cell pellet resuspended in 1200 µL media and seeded into each well of n-well plates or µ-Slide with dilution factor **F** and incubated for 16 hours at 37°C in 5% CO2.

**Confocal Microscopy**

Around $1*10^4$ cells are seeded into each well of 8-well µ-Slide (F ~ 300) and incubated overnight.

Transfection mix of 25 µL Opti-MEM I containing 0.3 µL MessengerMAX and 100 ng mock treated / acylated EGFP mRNA was prepared according to manufacturer instructions. After washing each well with 1X PBS, cell was supplied with 175 µL Opti-MEM I and the mix was added dropwise to the cells and transfection was incubated for 6 h at 37°C. Media was removed, and cells were washed with 1X PBS. For RNA release, denoted concentration of A + B (4:1) was mixed well in 1X PBS and added to the wells respectively, while blank treatment of 1X PBS was applied to other wells, and cells were incubated at 37°C for 2 h. Cells were then washed thoroughly with 1X PBS twice and supplied with 200 µL fresh media and incubated for another 18 h. Media was removed, cell resupplied with 200 µL fresh media containing 1 µg/mL Hoechst 33342 for nucleus staining for 20 min each well and proceeded for imaging.

**Flow Cytometry**

Around 1*$10^5$ cells are seeded into each well of 24-well plate (F ~ 30) and incubated overnight.

Transfection mix of 100 µL Opti-MEM I containing 1 µL MessengerMAX and 400 ng mock treated / acylated EGFP mRNA was prepared according to manufacturer instructions. After washing each well with 1X PBS, cell was supplied with 900 µL Opti-MEM I and the mix was added dropwise to the cells and transfection was incubated for 6 h at 37°C. Media was removed, and cells were washed with 1X PBS. For RNA release, 4 mM A + 1 mM B (4:1) was mixed well in 1X PBS and added to the wells respectively, while blank treatment of 1X PBS was applied to other wells, and cells were incubated at 37°C for 2 h. Cells were then washed thoroughly with 1X PBS twice and supplied with 1 mL fresh media and incubated for another 18 h. Media was removed, cell washed briefly with 1X PBS and added 0.3 mL Trypsin-EDTA and cells were incubated for 5 min at 37 degrees. Then the solution was neutralized with 0.5 mL FBS-containing media, cells suspended and proceed directly to flow cytometer for fluorescence quantification. Cell numbers each dataset: ~ 10000 entries.

**RNA interference assay**

Around 4*$10^5$ cells are seeded into each well of 12-well plate (F ~ 8) and incubated overnight.

Transfection mix of 100 µL Opti-MEM I containing 2.5 µL Lipofectamine 3000 and 1000 ng mock treated / acylated shPD-L1 (scramble shPD-L1 for NC) was prepared according to manufacturer instructions. After washing each well with 1X PBS, cell was supplied with 900 µL Opti-MEM I and the mix was added dropwise to the cells and transfection was incubated for 6 h at 37°C. Media was removed, and cells were washed with 1X PBS. For RNA release, 4 mM A + 1 mM B (4:1) was mixed well in 1X PBS and added to the wells respectively, while blank treatment of 1X PBS was applied to other wells, and cells were incubated at 37°C for 2 h. Cells were then washed thoroughly with 1X PBS twice and supplied with 1 mL fresh media and incubated for another 40 h. Transfected cells were harvested and lysed in RIPA buffer (protease added). After sonication, the cell lysates were analyzed in 12% polyacrylamide-SDS gels. Proteins on SDS-PAGE gel were transferred to a PVDF membrane, which was blocked with a 5% blocking grade milk powder-containing TBST buffer and then incubated with anti-PD-L1 monoclonal antibody (Proteintech) with a dilution factor of 1:2500. The primary antibody-bound blot was washed with 0.1% TBST buffer and then incubated with a secondary antibody (conjugated to AlexaFlour488, Thermo Fisher Scientific) with a dilution factor of 1:10000, washed with 0.1% TBST buffer and visualized on GE Amersham Typhoon 9410 fluorescence gel scanner.

#### Supplementary Figures

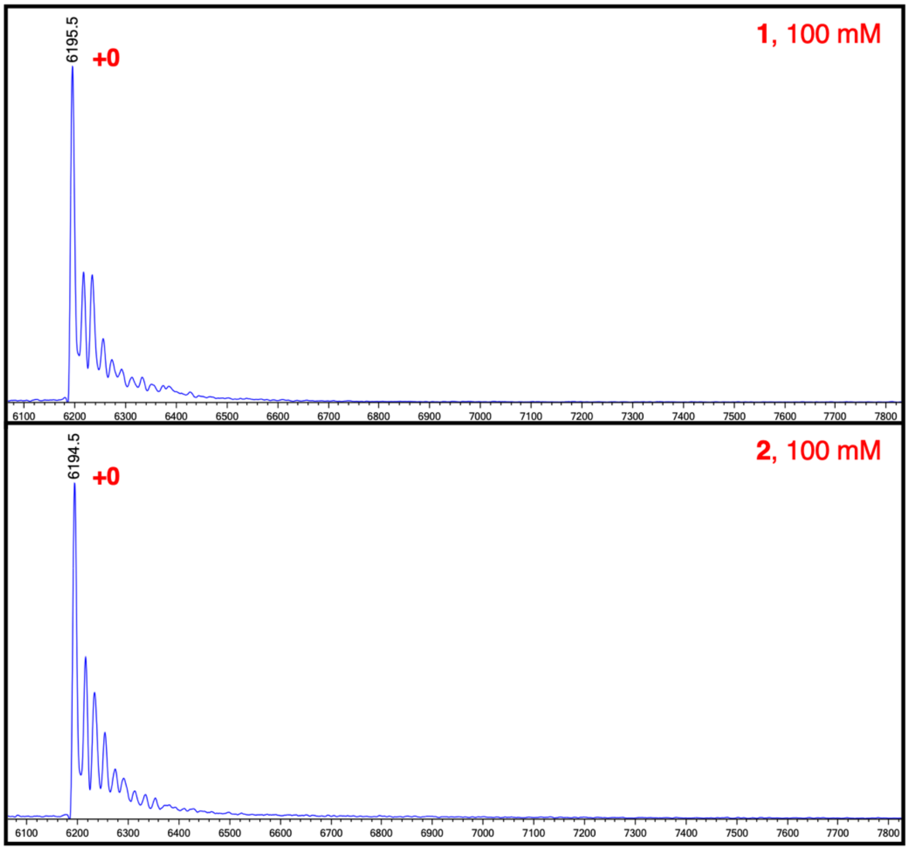


**Figure S 1. MALDI-TOF analysis of 1 and 2 treated 18-mer FAM-RNA. No reaction could be observed.**

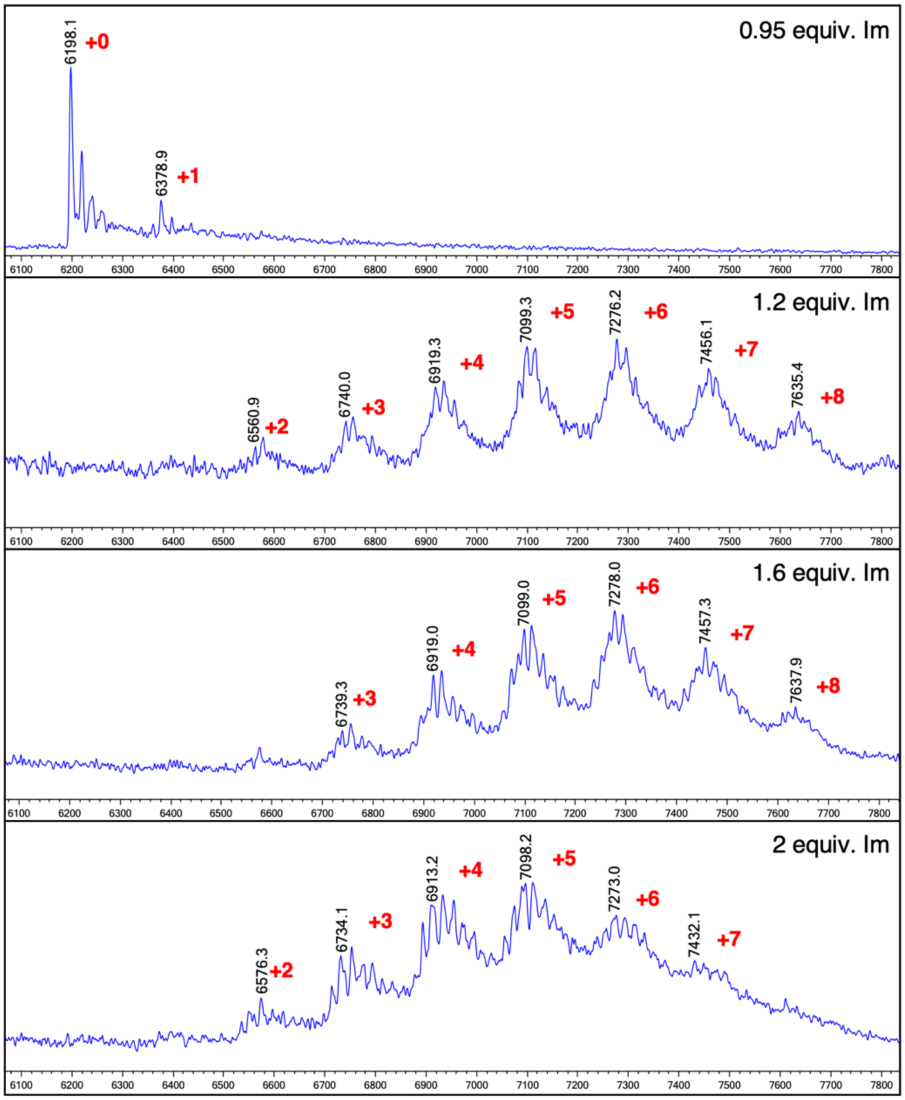


**Figure S 2. MALDI-TOF analysis of 18-mer FAM-RNA treated with PIC made from different ratio of 3 and 4. Im, Imidazole-2-amide (4). Equivalence was calculated based on 4-nitrobenzyl chloroformate (3).**

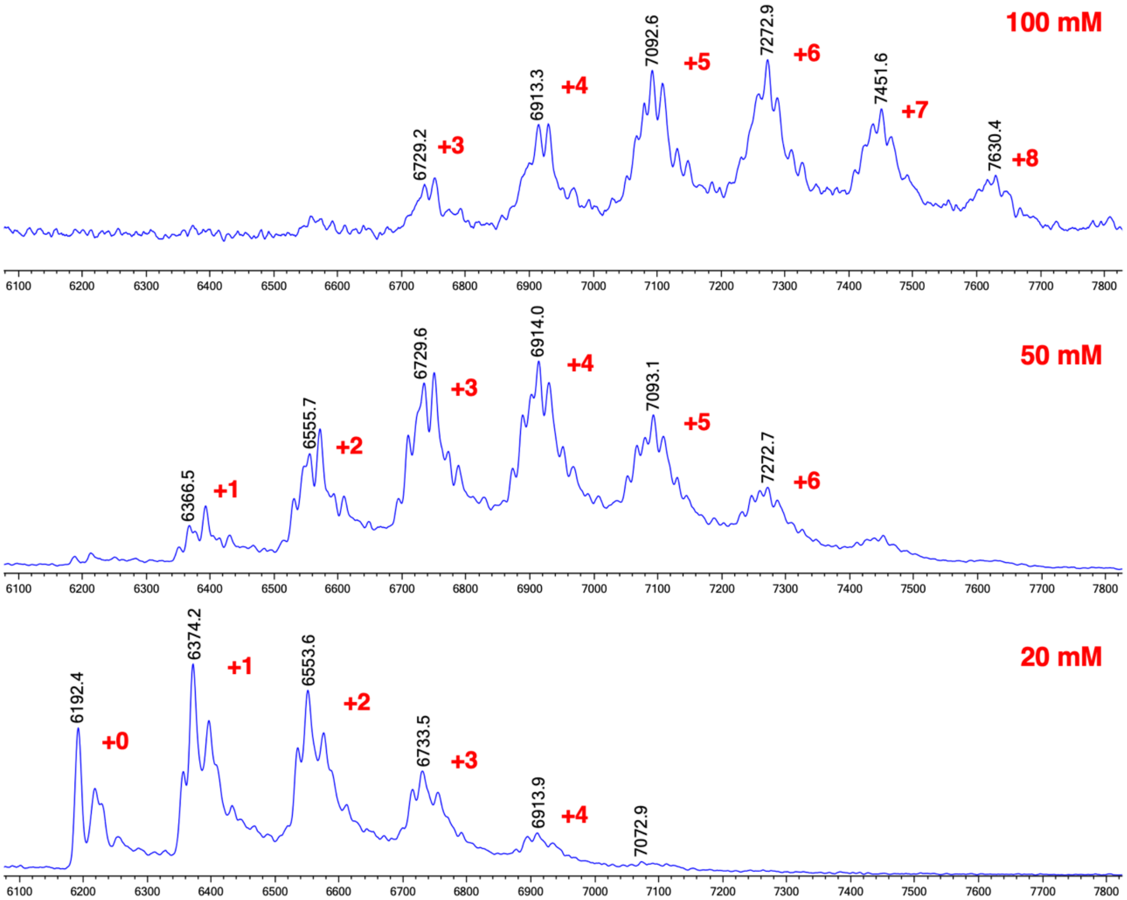


**Figure S 3. MALDI-TOF analysis of PIC concentration (20, 50, 100 mM) impact on 18-mer FAM-RNA modification.**

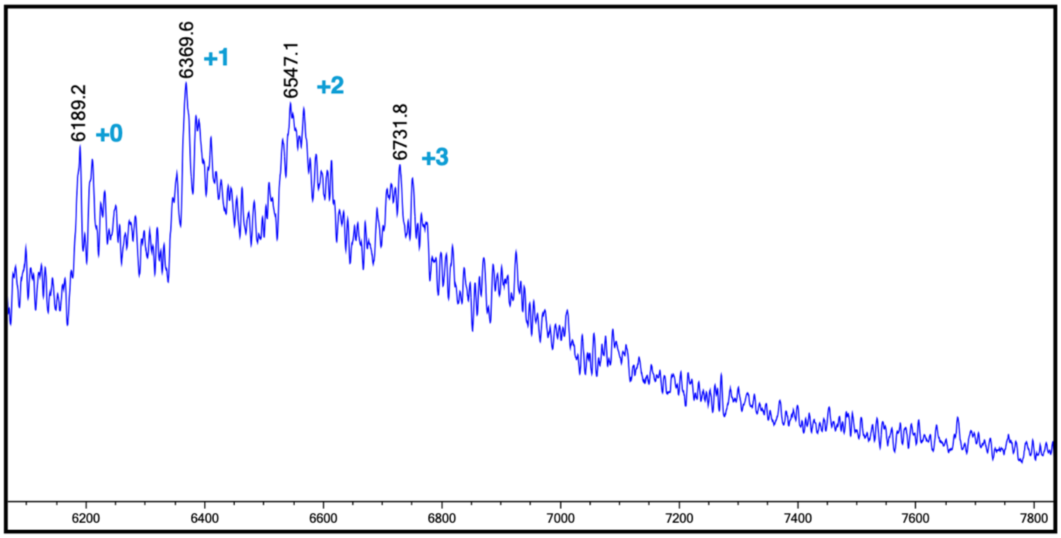


**Figure S 4. High-concentration nitroreductase treatment (500 ug/mL NTR, 5 mM NADH) of 100 mM PIC-acylated 18-mer RNA in 1X PBS at 37 °C for 24 h could not fully convert modified RNA to its original state (analyzed by MALDI-TOF).**

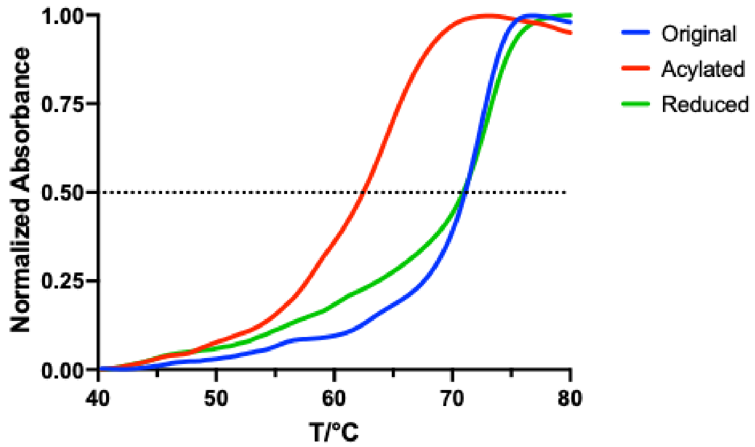


**Figure S 5. Melting point analysis of 39-mer original, acylated and reduced RNA oligo with full complementary DNA shows similar RNA-DNA duplex stability among reduced (Tm = 71.0°C) and original group (Tm = 71.2°C) with significantly destabilized duplex of acylated group (Tm = 62.5°C), proving hindrance in RNA base pair recognition when acylated, which is reversible by reduction. Melting curve of each group was normalized to 1.0 at maxima and 0.0 at minima. Tm was calculated based on interception of normalized curve with y = 0.5.**

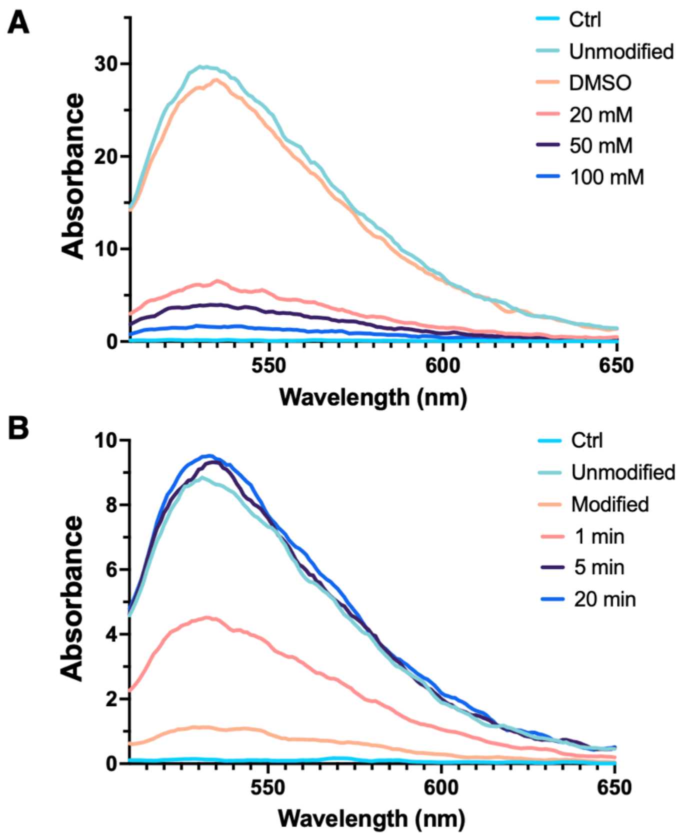


**Figure S 6. Fluorescence spectrum of equal amount of F30-pepper aptamer applied with control-recovery system. A) F30-pepper aptamer treated with different concentration of PIC. Ctrl, HBC530 only. DMSO, mock treatment of 20% DMSO. B) Kinetics of acylated F30-pepper release with THDB-BIPY. Modified, 50 mM PIC acylated aptamer.**

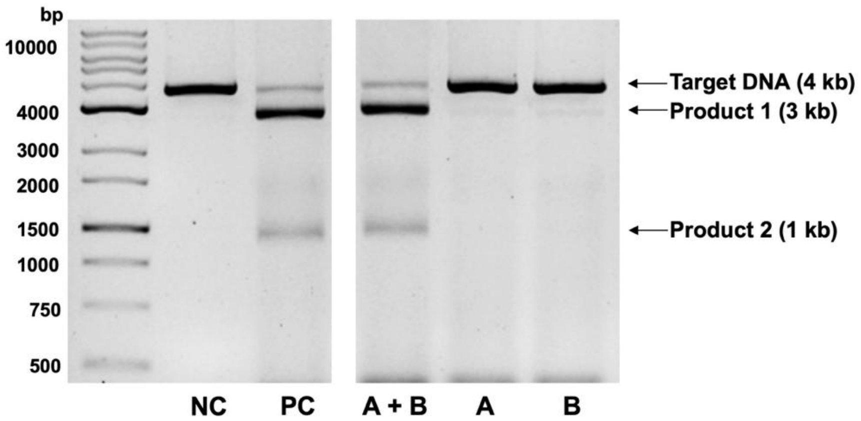


**Figure S 7. Treatment of A only (4 mM) and B only (1 mM) on modified sgRNA could not trigger functional restoration and Cas9 cleavage of target, analyzed on agarose gel electrophoresis. A + B (4 / 1 mM) was included as reference. NC, target DNA only. PC, target DNA treated with unmodified sgRNA and Cas9.**

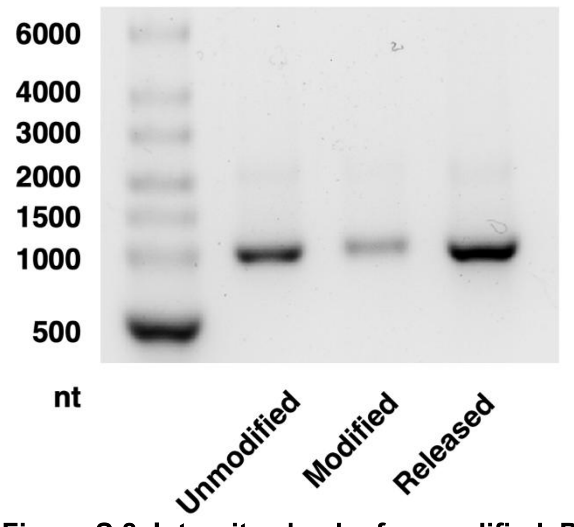


**Figure S 8. Integrity check of unmodified, PIC (35 mM) modified and THDB-BIPY (4 / 1 mM) released mEGFP on 1% agarose gel showing no smear band or sign of truncation, indicating good integrity.**

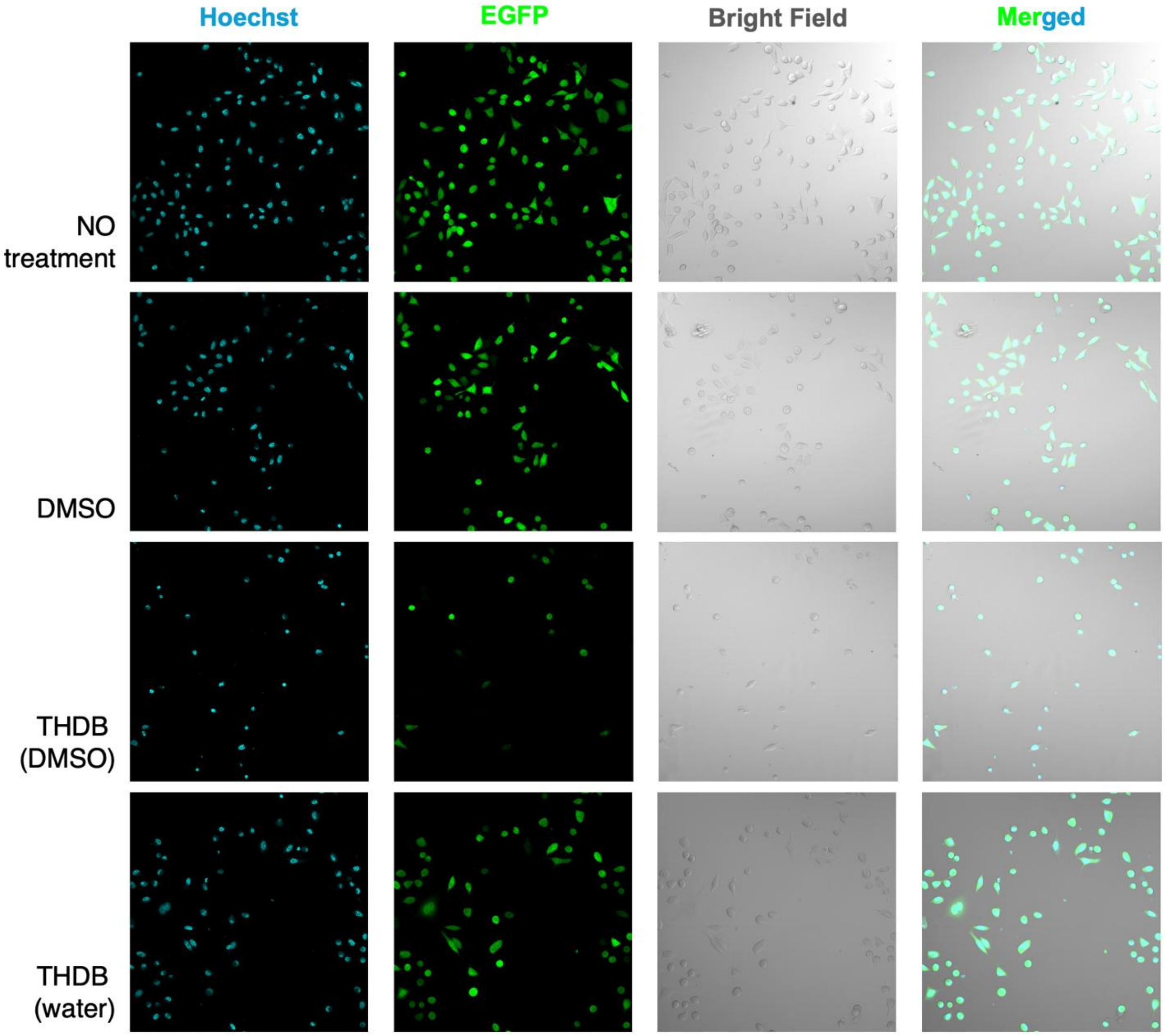


**Figure S 9. Comparison of mEGFP-transfected HeLa cells treated with DMSO, aqueous THDB stock and THDB stock in DMSO with microscopy shows that THDB agent prepared in DMSO drastically inhibit the translation of mEGFP and causes a decrease in cell numbers compared to all other entries (bright field scanned to show changes in cell density). Data are representatives of two replicate experiments.**

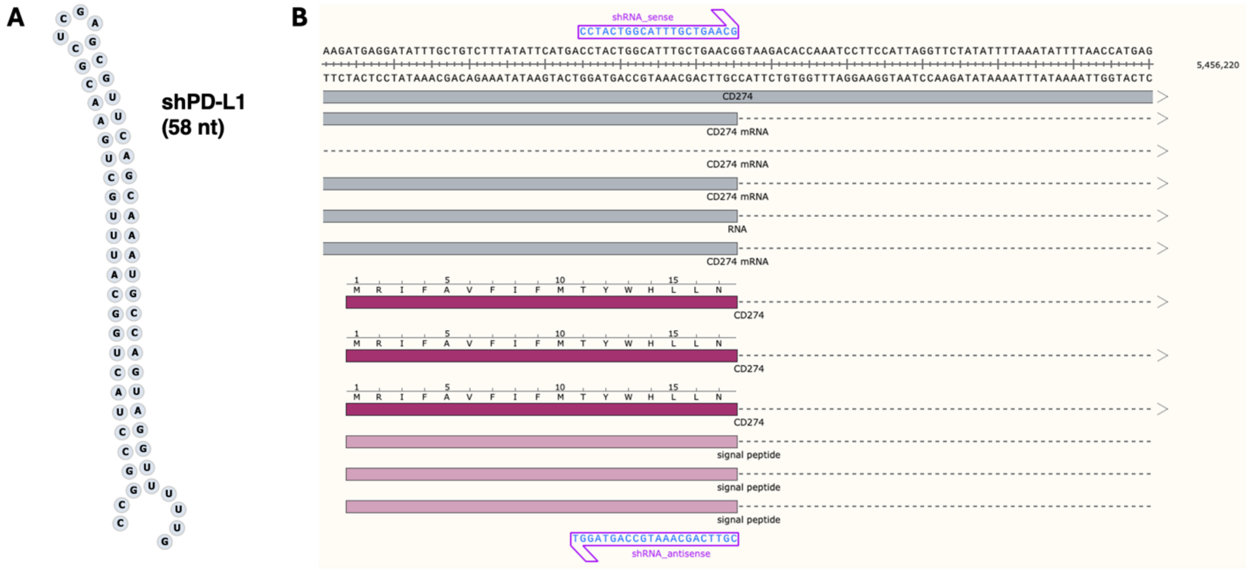


**Figure S 10. Design of short hairpin RNA targeting PD-L1 (CD274). A) Sequence of designed shPD-L1 containing sense and antisense strands, conserved loop and overhang. B) Targeting region of shPD-L1 in human cell expression of PD-L1.**

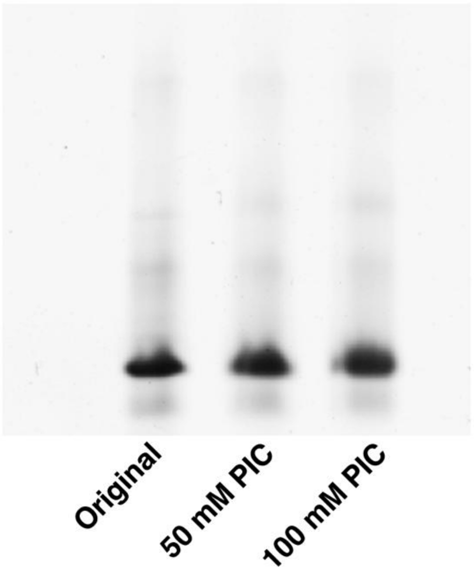


**Figure S 11. Integrity check of shPD-L1 after acylation by PAGE shows no change of RNA quality. The shRNA was modified directly after purchased without further purification.**

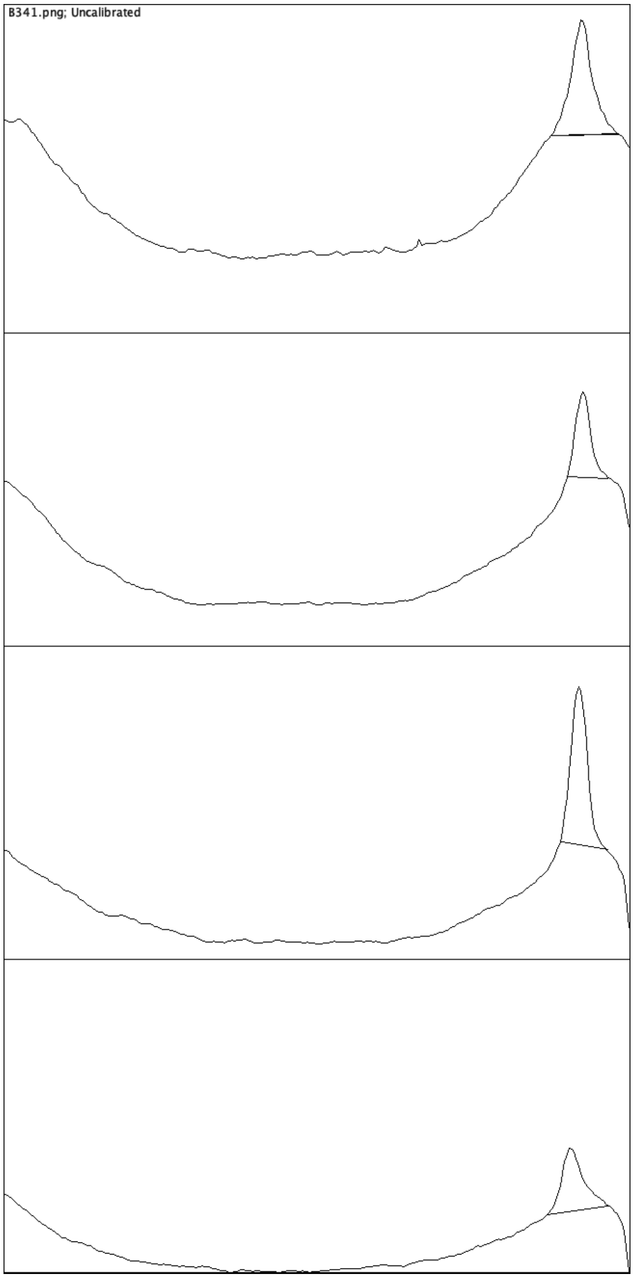


**Figure S 12. Integration of PD-L1 band intensity of each separate lane (NC, PC, Ac, SM) on immunoblot (See Figure 6C).**

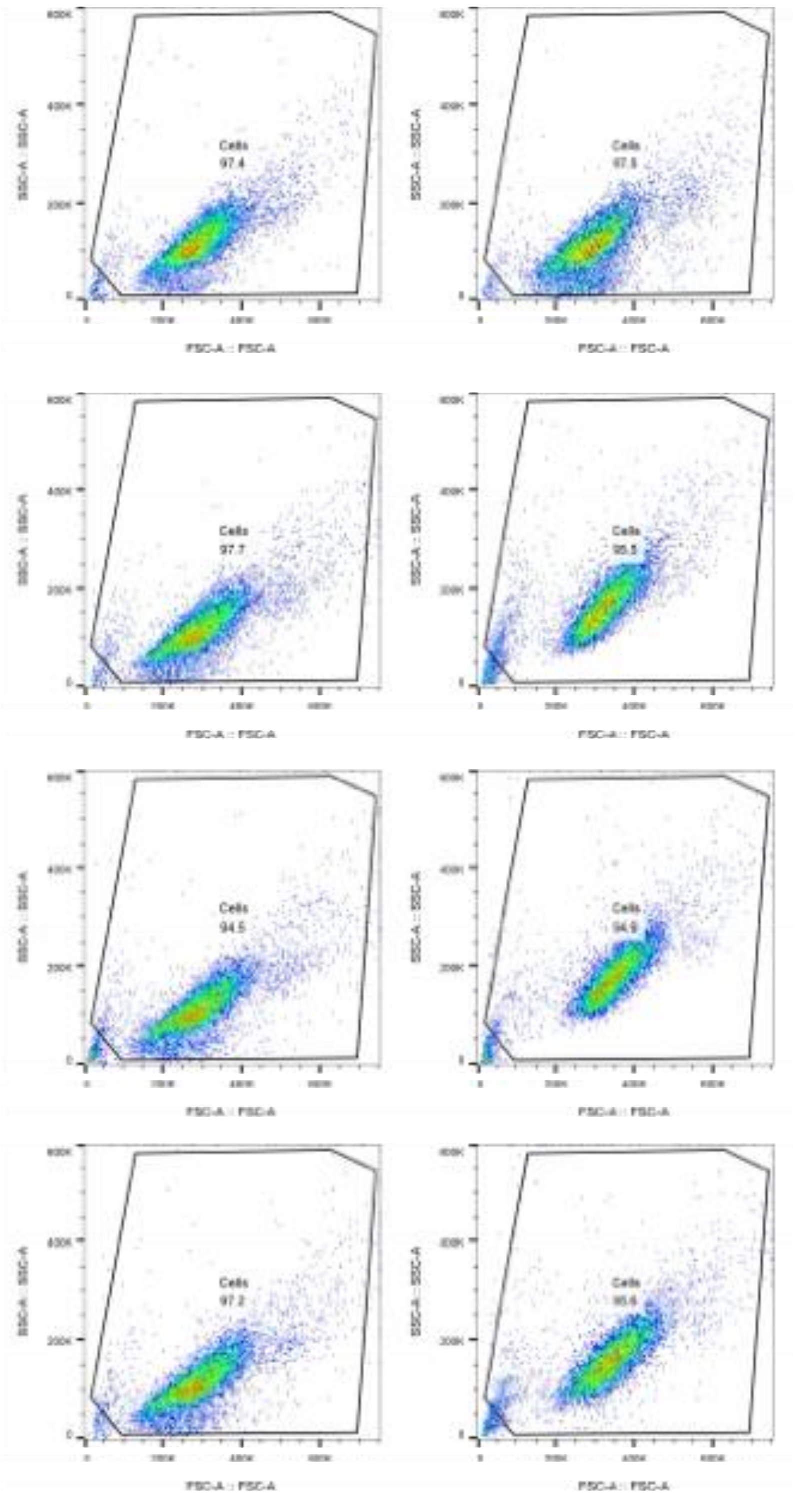


**Figure S 13. Flow cytometry cell gating. SSC: Side scatter; FSC: Forward scatter. A: Area.**

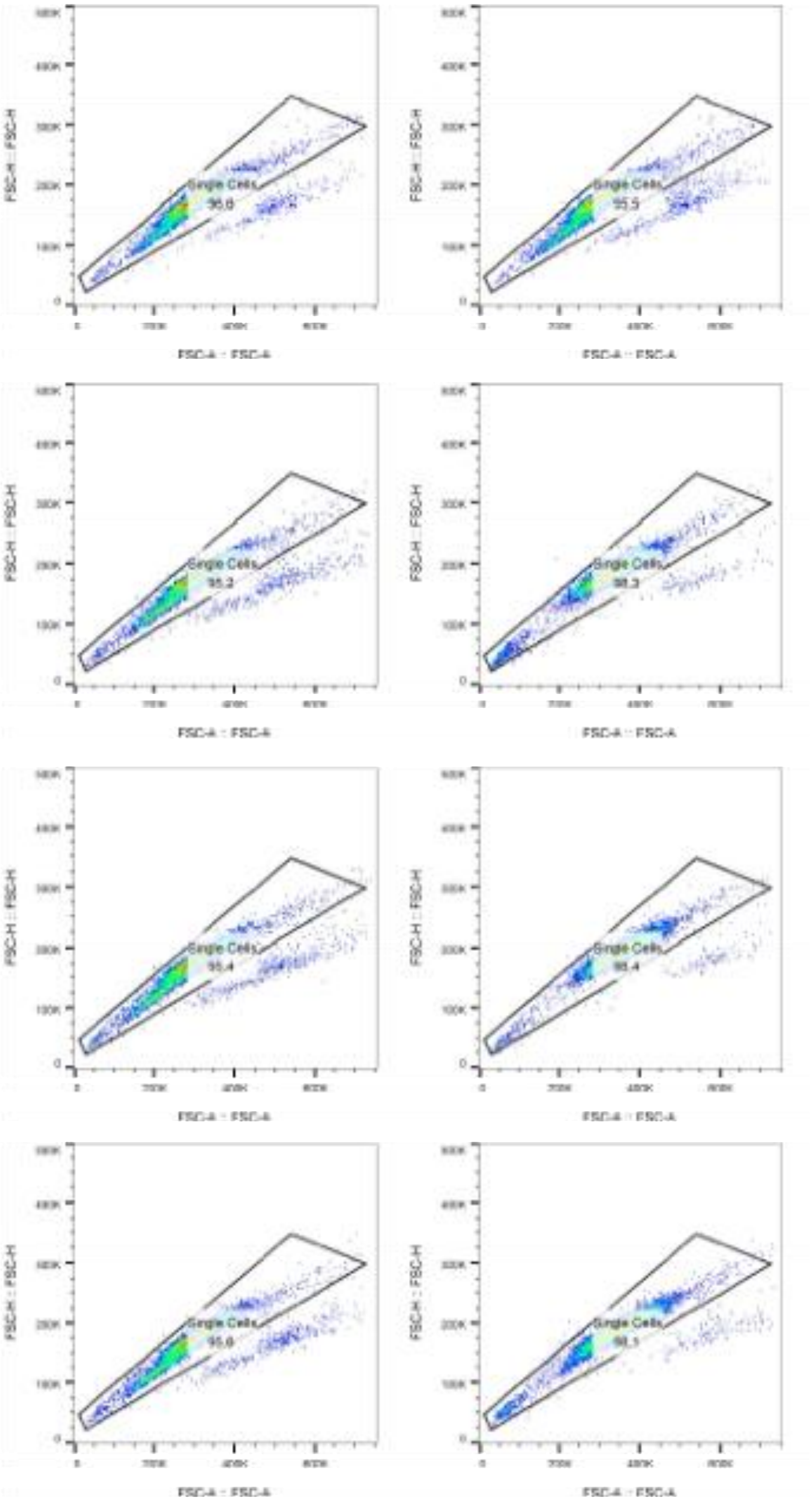


**Figure S 14. Flow cytometry single cell gating. FSC: Forward scatter. H: Height. A: Area.**

**Full Gel and Membrane:

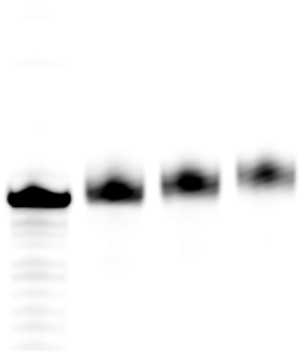

**Figure S 15. Full uncropped gel of Fig 1F.**

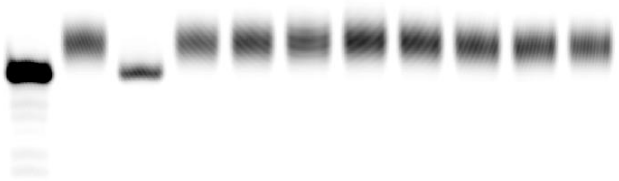

**Figure S 16. Full uncropped gel of Fig 2E.**

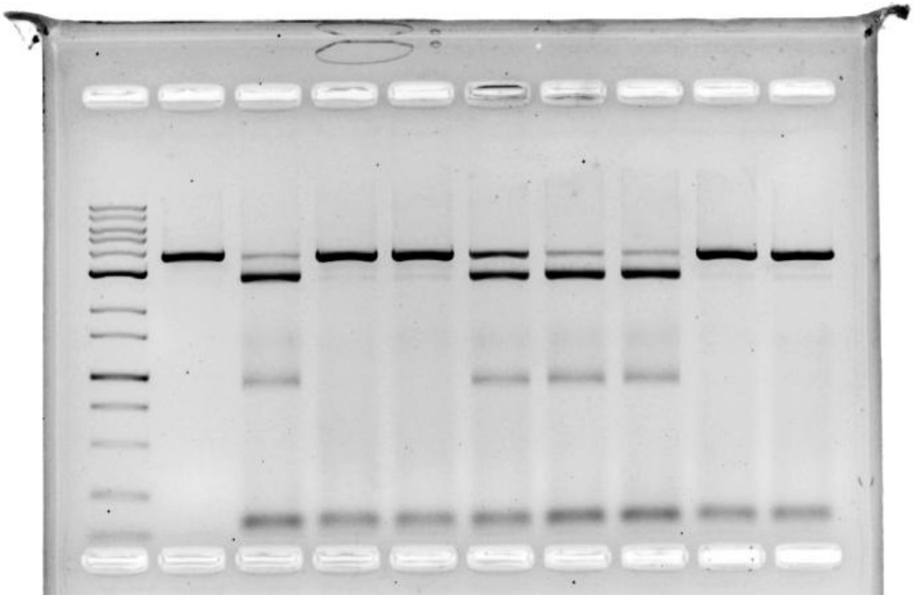

**Figure S 17. Full uncropped gel of Fig 4C and Fig S7.**

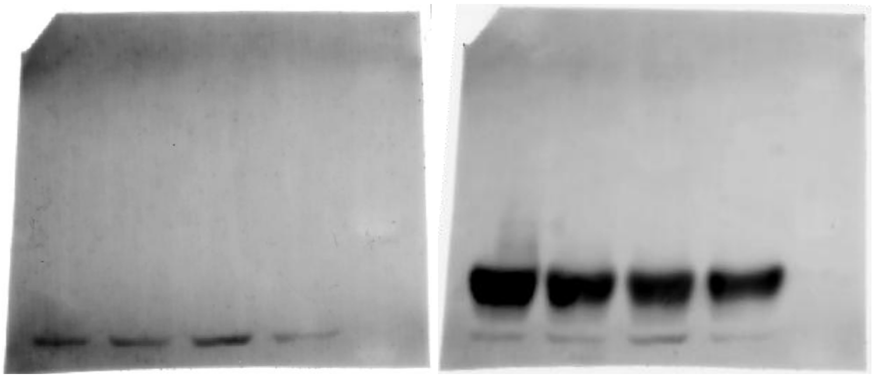

**Figure S 18. Full membrane of Western Blotting (after target & housekeeping monoclonal antibody incubation, respectively).**

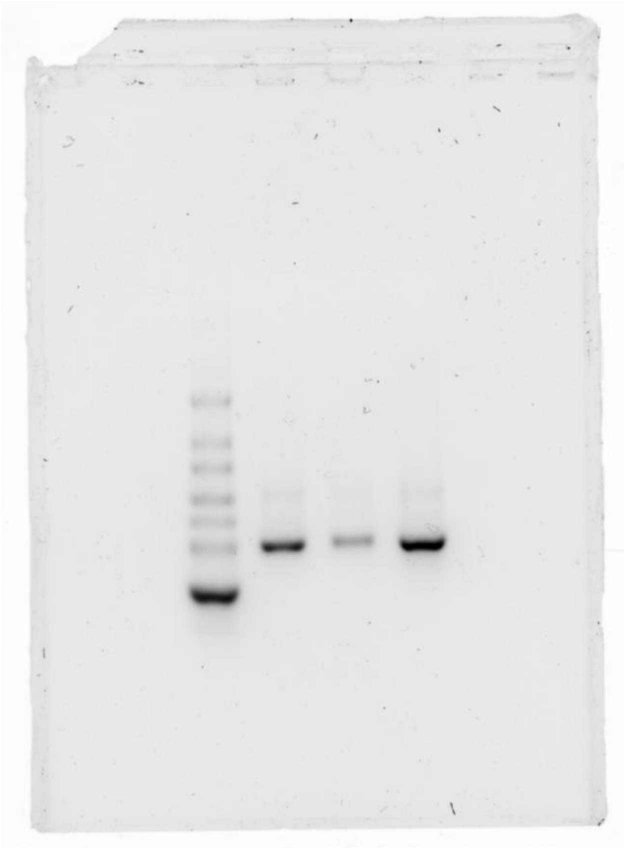
Figure S 19. Full uncropped gel of Fig S8.

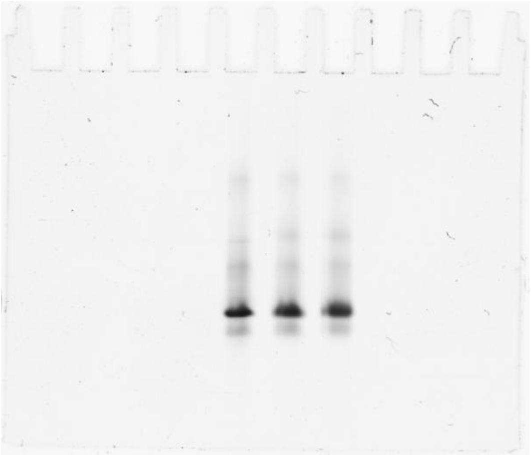
Figure S 20. Full uncropped gel of Fig S11.

### NMR Spectroscopy and Mass Spectrometry

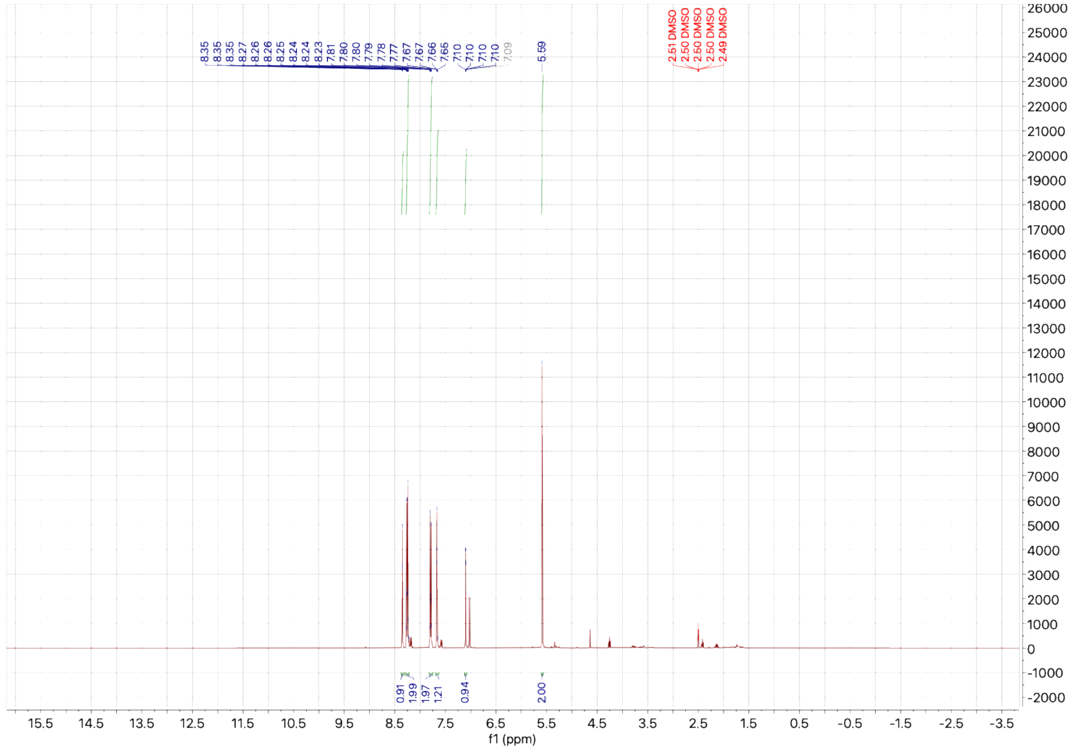


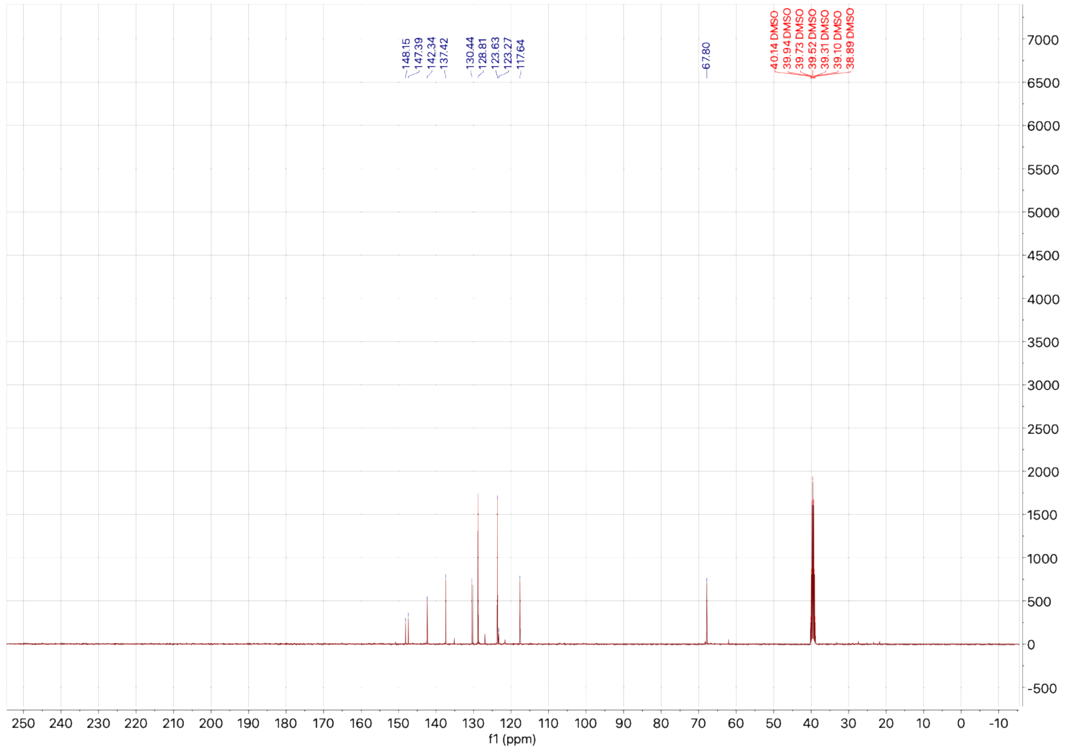


**Figure S 21. 1H and 13C NMR spectrum of 1.**

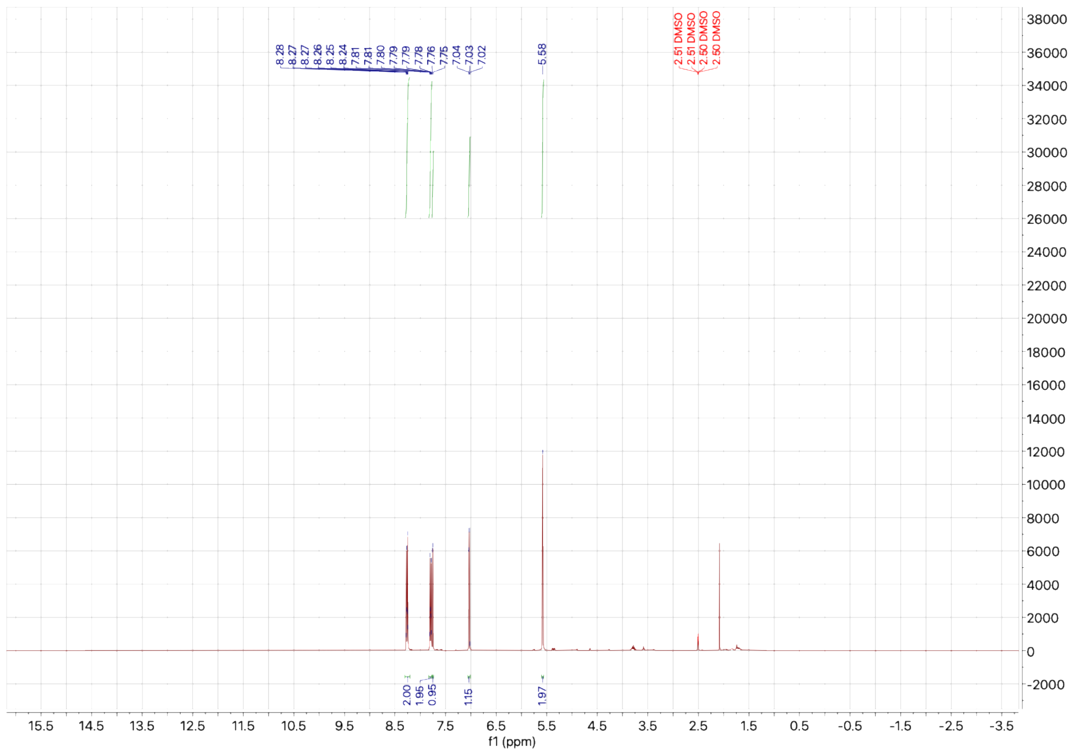


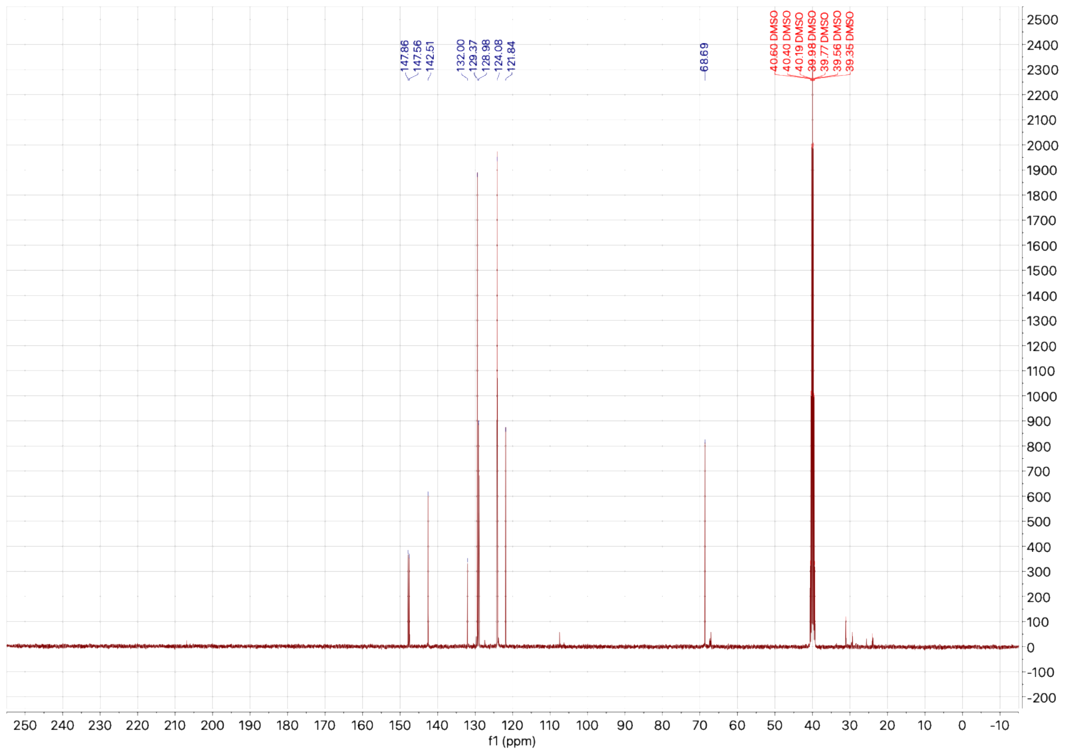


**Figure S 22. 1H and 13C NMR spectra of 2.**

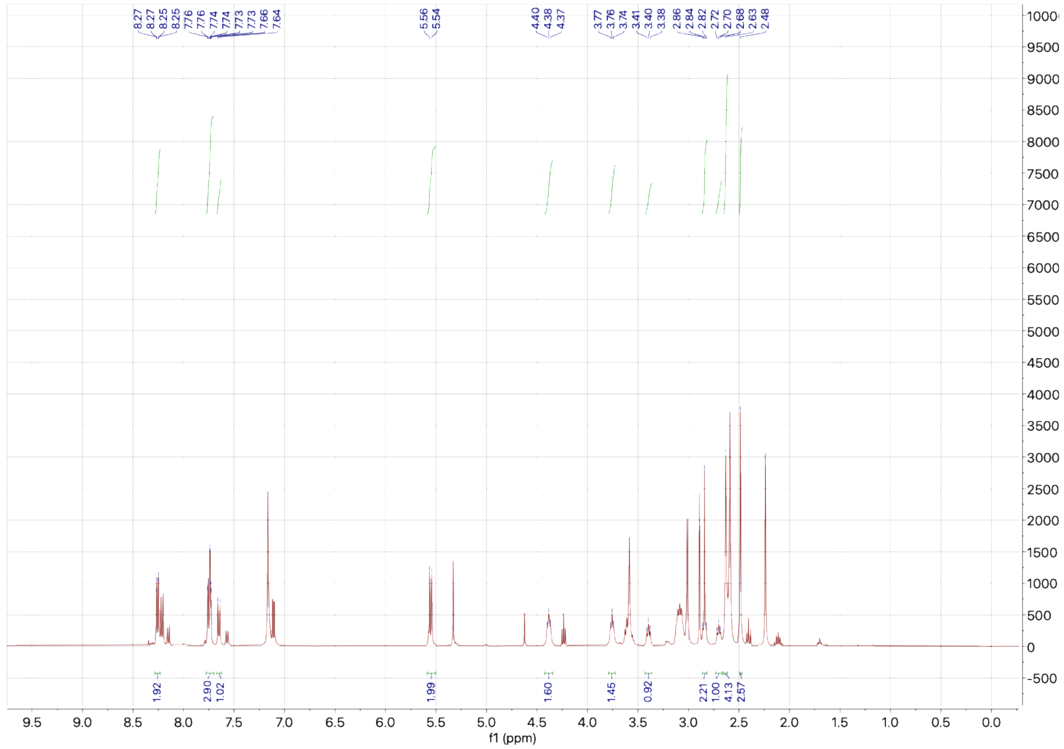


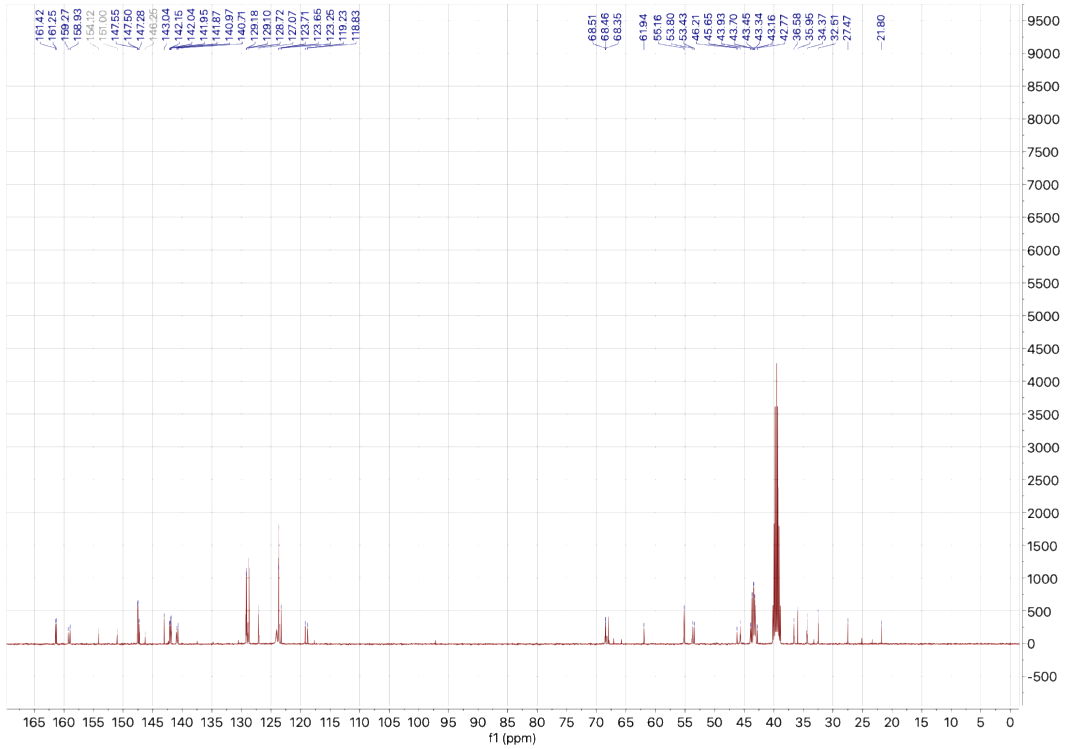


**Figure S 23. 1H and 13C NMR spectra of PIC.**

| Meas. m/z | # | Formula | Calc. Mass | Err [ppm] |
|---|---|---|---|---|
| 248.0659 | 1 | C11 H10 N3 O4 | 248.0666 | 2.82 |

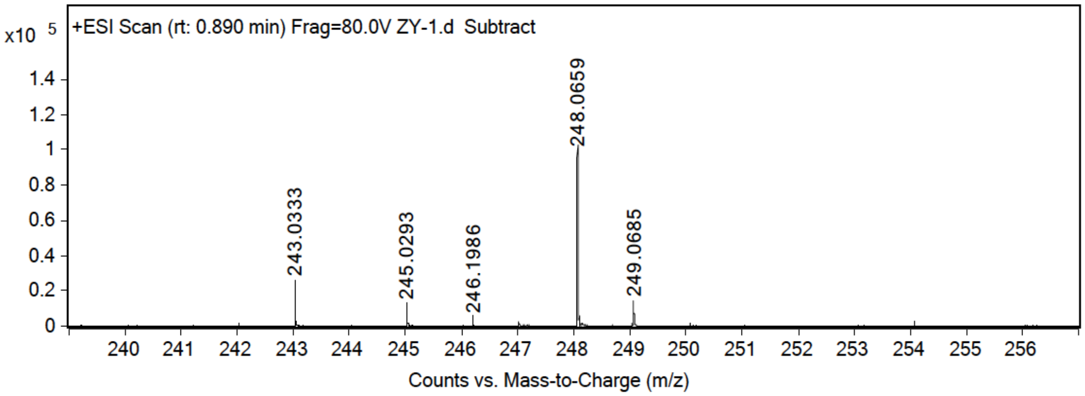


| Meas. m/z | # | Formula | Calc. Mass | Err [ppm] |
|---|---|---|---|---|
| 282.0271 | 1 | C11 H9 Cl N3 O4 | 282.0276 | 1.77 |

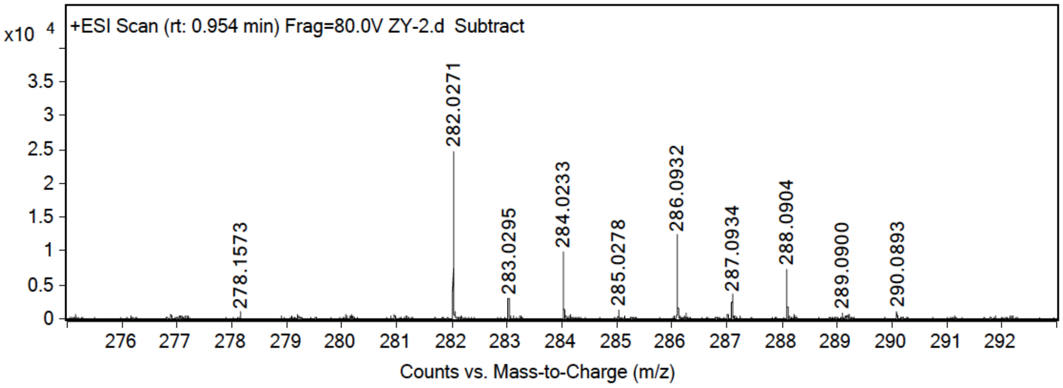


| Meas. m/z | # | Formula | Calc. Mass | Err [ppm] |
|---|---|---|---|---|
| 376.1616 | 1 | C17 H22 N5 O5 | 376.1615 | 0.27 |

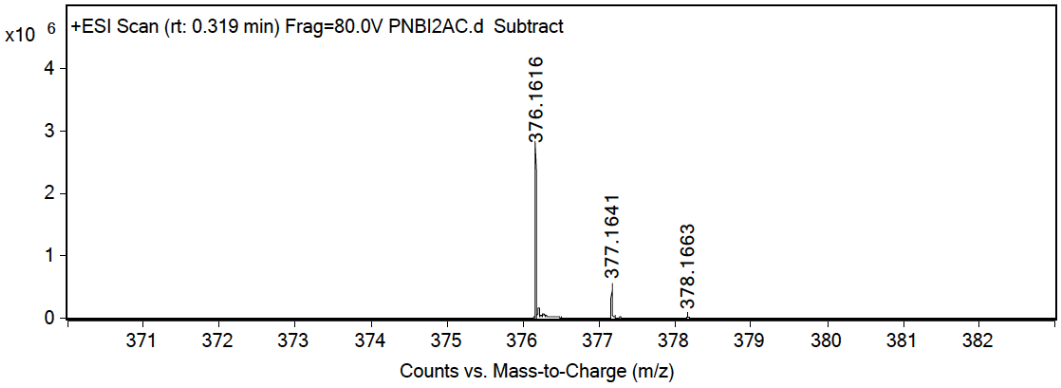


**Figure S 24. HR-MS analysis of 1, 2 and PIC.**

**REFERENCE**

(1) *GPP Web Portal - Design Hairpins to Target a Transcript Sequence*. broadinstitute.org, https://portals.broadinstitute.org/gpp/public/seq/search (accessed 2025-10-15).

(2) Velema, W. A.; Kool, E. T. Water-Soluble Leaving Group Enables Hydrophobic Functionalization of RNA. *Org Lett* **2018**, *20* (20), 6587-6590. DOI: 10.1021/acs.orglett.8b02938.
